# The majority of microorganisms in gas hydrate-bearing subseafloor sediments ferment macromolecules

**DOI:** 10.1101/2022.05.19.492412

**Authors:** Chuwen Zhang, Yun-Xin Fang, Xiuran Yin, Hongfei Lai, Zenggui Kuang, Tianxueyu Zhang, Xiang-Po Xu, Gunter Wegener, Jiang-Hai Wang, Xiyang Dong

## Abstract

**Background:** Gas hydrate-bearing subseafloor sediments harbor a large number of microorganisms. Sedimentary organic matter and upward methane fluids represent two important sources of carbon and energy for deep biosphere. However, which metabolism dominates the deep subseafloor of gas hydrate zone is poorly constrained. Here we studied the microbial communities in gas-hydrate rich sediments up to 49 meters below seafloor recovered by drilling in the South China Sea. We focused on distinct geochemical conditions, and performed metagenomic and metatranscriptomic analyses to characterize microbial diversity and their role in carbon mineralization.

**Results:** Comparative microbial community analysis revealed that samples above and in sulfate-methane interface (SMI) zones clearly distinguish from those below the SMI. Chloroflexota, are most abundant above the SMI, whereas Caldatribacteriota dominate below the SMI. Verrucomicrobiota, *Bathyarchaeia* and Hadarchaeota were similarly present in both types of sediment. The genomic inventory and transcriptional activity suggest roles in fermentation of macromolecule. By contrast, sulfate reducers and methanogens, organisms that catalyze the consumption or production of commonly observed chemical compounds in sediments are rare. Methanotrophs of the ANME-1 group thrived in or slightly below the current sulfate methane interface. Rare members from *Heimdallarchaeia* were identified to encode the potential for anaerobic oxidation of short-chain hydrocarbons.

**Conclusions:** These findings indicate that fermentation of macromolecules is the predominant energy source for microorganisms in deep subseafloor sediments that are experiencing upward methane fluxes.

## Background

Gas hydrates, ice-like crystalline solids composed of water and hydrocarbons, are widely discovered in the deep subseafloor of every continental margin [1], e.g. at 300 meters below the seafloor (mbsf) in Forearc Basin [2], 45 to 125 mbsf in Hydrate Ridge (offshore Oregon, ODP Site 1244) [3], and 271 to 330 mbsf in the Nankai Trough [4]. Ecological studies based on single marker genes revealed abundant and diversified members of archaea and bacteria in the deep subsurface sediments associated with gas hydrates over the global ocean [2-6]. All known forms of life require sources of carbon and energy to thrive. The deep subseafloor ecosystem is highly energy-limited, and microbial reaction rates are amongst the lowest known on Earth [7, 8]. It remains elusive how the microorganism acquire carbon and energy sources in deep subseafloor sediments from the gas hydrate zone.

Marine sediments contain Earth’s largest pool of organic carbon, derived from primary production in the overlying water column, land-derived inputs, cell debris after the lysis and death, or exudates [9, 10]. Based on studies in various oceanic regions e.g. Guaymas Basin [11, 12], Eastern Gulf of Mexico [13] and the Helgoland mud area [14, 15], the vast majority of microbial community inhabiting surface sediments are proposed to be heterotrophs, utilizing organic matter to meet their carbon and energy demands. With the continuous sedimentation over the geologic time, the deposited organic matter is buried deeper and deeper under seafloor and becomes increasingly recalcitrant for microbial unitization [7, 14]. Yet, it has been proposed that deeply buried communities continue to live on the organic remains [16]. The sedimentary organic matter mainly consists of biological macromolecules including carbohydrates, lipids, proteins and nucleic acids as well as other complex substances like humic and fulvic acids [9, 17, 18]. Generally, the complex macromolecules first need to be broken down into oligomers and monomers for fermentation to smaller molecules, which feed into respiration or methanogenesis. The biogenic methane gas can be an important source for gas hydrate formation and accumulation [1, 19, 20].Subseafloor gas hydrates are vulnerable to dissociation under changing environmental conditions, e.g. the rising ocean temperature or dropping hydrostatic pressure, leading to the emission of gases (mostly methane) to the ocean and the atmosphere [21, 22]. In contact with sulfate these gases supplied from deep reservoirs serve as alternative energy and carbon sources for benthic microbes. Within sulfate methane interfaces, methane is consumed by anaerobic methane oxidizing archaea (ANME), typically forming syntrophic consortia with sulfate-reducing partner bacteria [23, 24]. In the surface or shallow sediments, methane fluxes were reported to greatly stimulate growth of ANMEs and coupled sulfate reducers, thus the deep biomass [6, 25]. However, it remains unclear whether the degradation of organic matter or methane fluid is the main metabolic strategy to sustain microbial life in deep subseafloor of gas hydrate zone.

In this study, we explored microbial community and their activities for carbon and energy acquirements in the deep subseafloor from the gas hydrate zone. Four cores were drilled from the gas hydrate zone in the South China Sea. Through the combination of geochemical measurements, metagenomics and metatranscriptomics, we provide evidence that unlike surface sediments, fermentative degradation of biological macromolecules is an important strategy to support deep biosphere experiencing methane fluxes.

## Results

### Geochemical characterization

Four cores from gas hydrate drilling sites (**Figure S1; see Methods**) were retrieved from the Shenhu area (*n* = 1; SH-W20A) and Qiongdongnan Basin (*n* = 3; QDN-W01B, W03B and W04B). All the four drilling sites were located above deep subsurface gas chimneys, indicating upward gas fluxes within sediments [26, 27]. All cores except QDN-W04B contained gas hydrates with various morphologies. The natural gases in the Shenhu area and Qiongdongnan Basin were reported to consist of predominantly methane accompanied by C_2_–C_5_ gases [28, 29]. Aligning with this, methane and non-methane gaseous alkanes including ethane and propane were detected in these four cores (**Table S1**).

Based on porewater sulfate and methane profiles plus the extrapolation of linear sulfate gradients [30-32], we predicted their depth distributions of the sulfate-methane interface (SMI; **Figure 1a**). For QDNB-W03B, QDNB-W04B, and SH-W20A, the SMI depths should be around 28, 40 and 30 mbsf, respectively [6, 32]. The DIC profiles also supported the active sulfate reduction above SMI, with the increased DIC concentrations above SMI and gradually decreased DIC contents after sulfate depletion (**Figure 1b**). In parallel, δ^13^C values of DIC were slightly negative at the top of the cores (−7.6∼−18.2‰) and became more depleted along with the increased sediment depth (−23.9∼−33.2‰). Such decrease can be caused by the oxidation of isotopically depleted methane. Below the depth of SMI, δ^13^C_DIC_ values increased to −11.6∼−13‰, indicating a dominance of organic matter degradation (**Figure 1b**). For these three cores, total organic carbon (TOC) contents in sediments decreased with depth (**Figure 1c**), suggesting active microbial carbon mineralization [25, 33].

**Figure 1.**
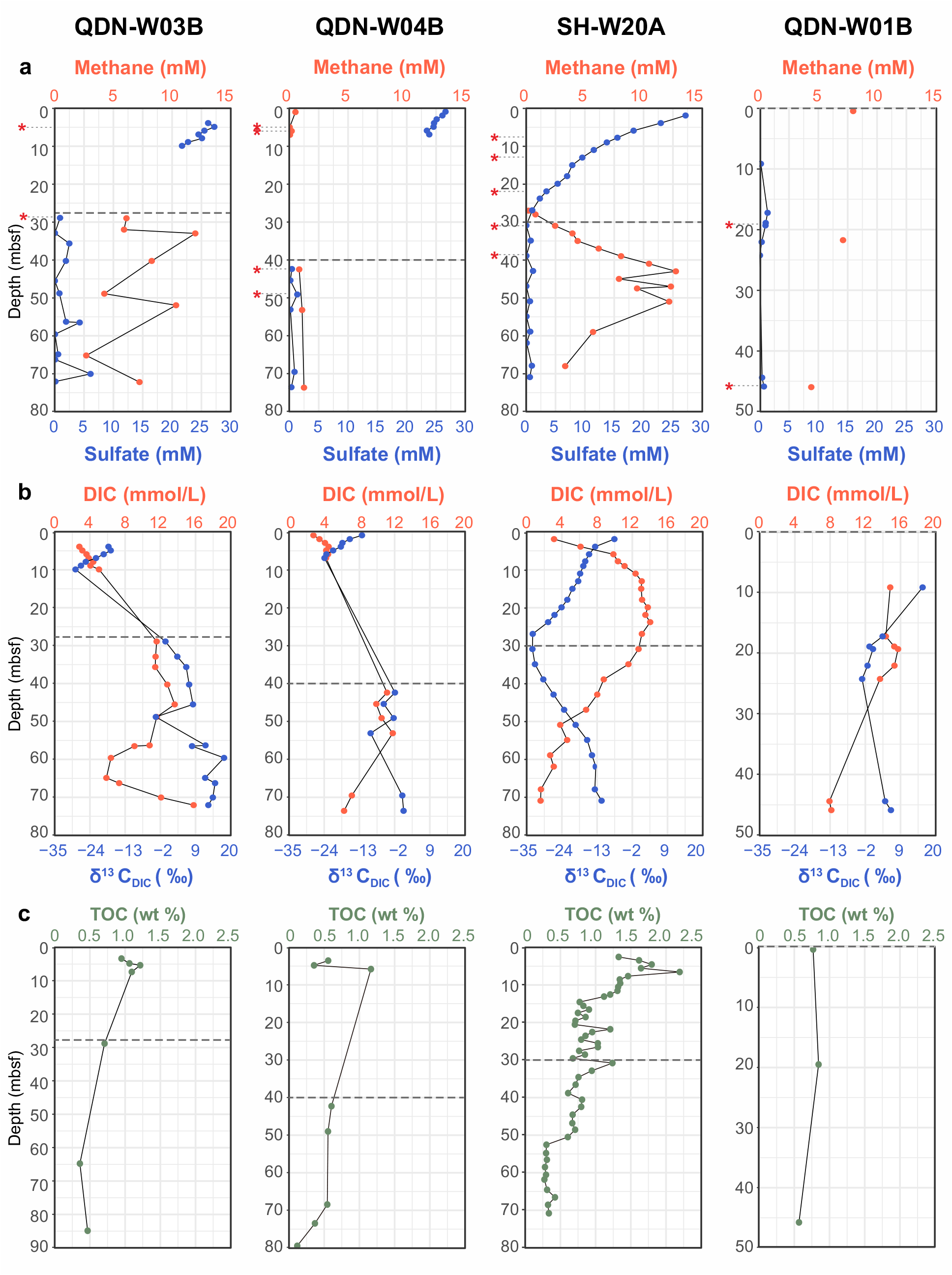
Sediment geochemistry of samples cored in the hydrate zone. Depth profiles of (a) methane and sulfate, (b) DIC content and δ^13^C values of DIC, and (c) TOC content. Dashed lines mark the predicted sulfate-methane interface predicted based on linear sulfate gradients. Samples used for metagenomic and metatranscriptomic analyses are marked with red asterisks.

In contrast, QDN-W01B shows low sulfate concentrations (<1.5 mM) in the entire sampled core (9−45 mbsf; **Figure 1a**), and concentration of methane were in the range of a few millimoles. These porewater profiles indicated a shallow SMI near the sediment/water interface [6, 32]. At 9−45 mbsf of sediments in this core, TOC contents were observed to range between 0.57−0.85% (**Figure 1c**). Simultaneously, DIC concentrations decreased, whereas δ^13^C_DIC_ values initially decreased and then increased with depth (**Figure 1b**). These data suggested the potential occurrence of microbial fermentation process at these depths considering the without availability of sulfate acceptor [33].

### Microbial diversity over redox zonation

To compare deep subseafloor microbiomes in response to distinct redox zones, 13 sediment samples were selected for shotgun metagenomic sequencing based on porewater sulfate concentrations and available DNA yields. For each sediment core, the alpha diversity analysis based on single-copy marker genes revealed a decline in both Chao1 and Shannon diversity with depth (**Figure 2a**). When grouping samples according to the biogeochemical zonation, subseafloor sediments above SMI supported significantly (Wilcoxon rank sum test, *P* < 0.001) more diverse communities than those below SMI (**Figure S2**). This is evidenced by higher values of Chao1 (966 ± 247 vs 315 ± 113) and Shannon (5.14 ± 0.32 vs 3.51 ± 0.59) above SMI (**Figure 2a**), highlighting the importance of sulfate availability in shaping microbial community compositions.

**Figure 2.**
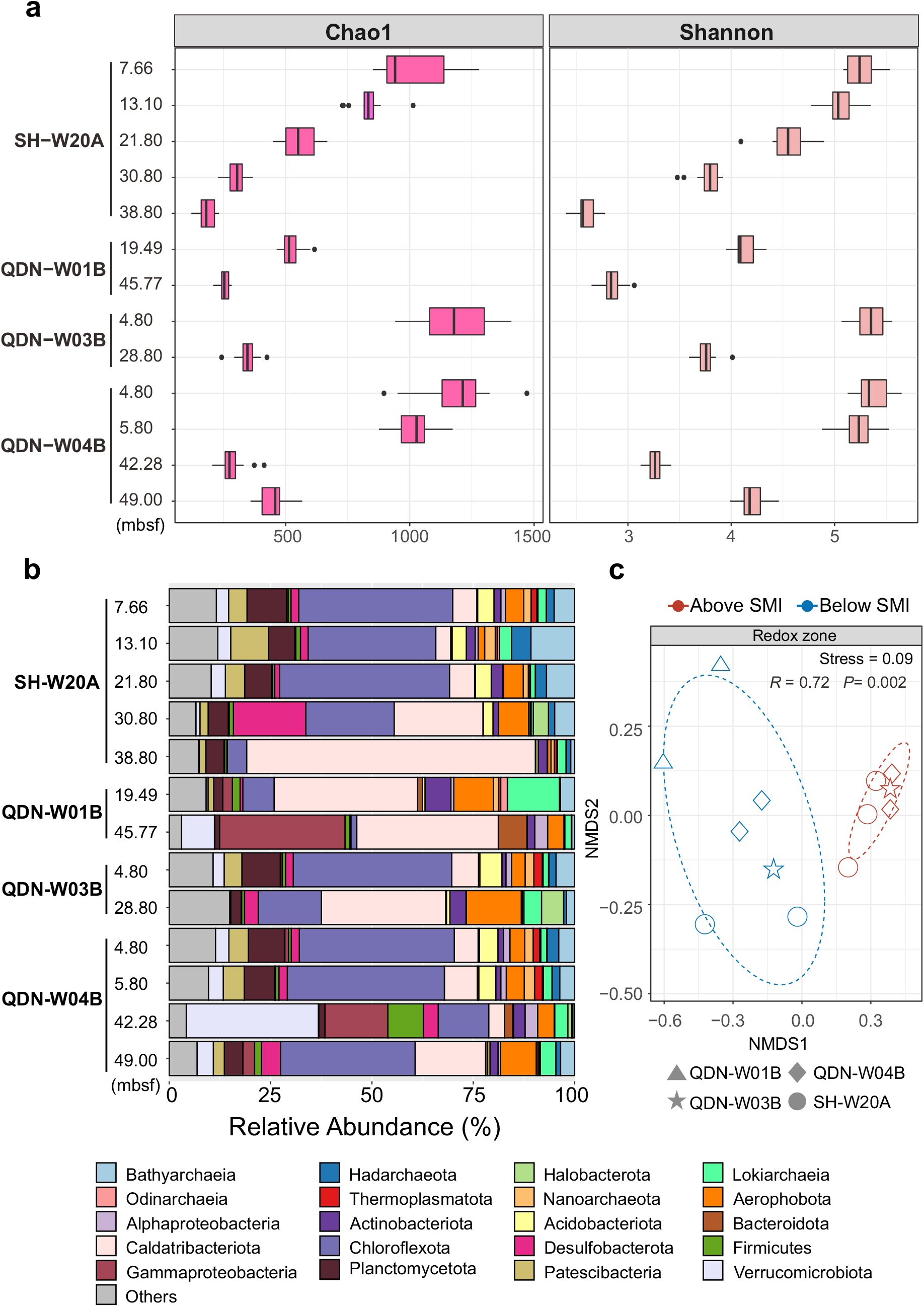
Microbial composition and diversity analysis of the deep subsurface sediments from the gas hydrate zone. (a) Alpha diversity of microbial communities based on metagenomic reads of 14 single-copy marker genes. (b) Taxonomic compositions of microbial communities based on 16S miTags extracted from the metagenomes. Full taxonomic information is provided in **Table S2**. (c) NMDS analysis of a Bray-Curtis dissimilarity matrix calculated from the single-copy marker gene *rplB* OTU table. Testing for significant differences in community structure between redox zones was performed using ANOSIM with 999 permutations.

For microbial community profiling, we classified 16S rRNA gene fragments (i.e., 16S miTags) from shotgun metagenomic reads (**Figure 2b**). In QDNB-W01B, Caldatribacteriota was the most abundant lineage (on average 35.2% of the whole community), followed by *Gammaproteobacteria* (16.7%), *Lokiarchaeia* (7.2%) and Aerophobota (6.9%). The microbial communities from QDNB-W01B were similar to those from the SMI-below sediments of SH-W20A, QDNB-W03B and QDNB-W04B (**Figure 2b**). For the SMI-above sediments of the latter group, Chloroflexota was the most abundant lineage (37.9%), followed by Planctomycetota (8.1%), *Bathyarchaeia* (5.8%), and Acidobacteriota (4.3%). The beta diversity analysis confirmed that deep subseafloor community variations strongly correlated with redox zones (ANOSIM, *R* = 0.72, *P* = 0.002; **Figure 2c**). Member of the Halobacterota phylum were observed to be only abundant in sediments below SMI from SH-W20A (3.7%) and QDN-WO3B (5.5%), consisting of mostly ANME-1 and *Methanosarcinia* (**Figure S3**).

### Dominant members harbor capabilities to ferment diverse macromolecules

Metagenomic assembly and binning yielded 578 metagenome-assembled genomes (MAGs, < 99% ANI) with > 50% completeness and < 10% contamination (**Table S3 and Figure S4**). They clustered into 349 bacterial and archaeal species-level clades, with most present less than 1% relative abundance across all samples (**Figure S5 and Table S4**). For utilization of biological macromolecules, we screened these MAGs for the presence of genes encoding for CAZymes, peptidases and nucleases. In total, we detected 21942 potential CAZymes out of 577 MAGs (**Tables S5)** and 46693 potential peptidases out of 578 MAGs, respectively (**Tables S6)**. This result suggested the utilization of biological macromolecules as general features for those subseafloor microorganisms in the gas hydrate zone. Approximately 2.3% CAZymes and 2.6% peptidases could be potentially released to the environment (**Tables S5 and S6**). The secretory CAZymes and peptidases are important for the cleavage of polymeric substrates before their incorporation into cells [34].

Of the 21942 CAZymes hits (**Figure S6**), genes belonging to glycosyltransferases were the most abundant (46%), followed by glycoside hydrolases (32%), carbohydrate esterases (9%) and carbohydrate-binding modules (5%). Carbohydrate degradation in the deep subseafloor of gas hydrate zone was carried out by members from phylogenetically diverse microbial phyla/classes, including dominant bacterial and archaeal lineages of Chloroflexota, Caldatribacteriota, Verrucomicrobiota, *Bathyarchaeia, Lokiarchaeia* and other phylogenetic clusters (**Figure 3 and Table S7)**. Most carbohydrate-degrading microbes contained diverse extracellular CAZymes allowing the cleavage of multiple carbohydrates and numerous sugar transport systems for oligo-/monomers uptake (**Figure S7**). Consequently, these microorganisms should be able to utilize a broad spectrum of carbohydrates including chitin, cellulose, pectin, polyphenolics, starch, xylans and xyloglucan (**Table S8**). The simple sugars produced by the activity of extracellular CAZymes can enter into the glycolysis pathway which was prevalent across bacterial and archaeal lineages (**Figure 3)**. Most of bacterial and archaeal MAGs lacked genes for respiration. They instead encoded the potential to metabolize pyruvate produced during glycolysis to acetyl-CoA and further into fermentation pathways (**Figure 3)**, yielding various organic acids. These acids mainly included acetate (*acdA* or *pta*+*ack*), formate (*fdoG* or *pflD*) and lactate (*ldh*). In addition, fermentative hydrogen production (Nife group 3, Nife group 4a-g) might function as another electron sink for anaerobic degradation of macromolecules. Gene-centric surveys also show genes encoding for formate (up to 163.18 genes per million, GPM), acetate (up to 158.81 GPM), lactate (up to 43.68 GPM) and hydrogen (up to 210.91 GPM) fermentation to be highly abundant (**Figure 4a**).

**Figure 3.**
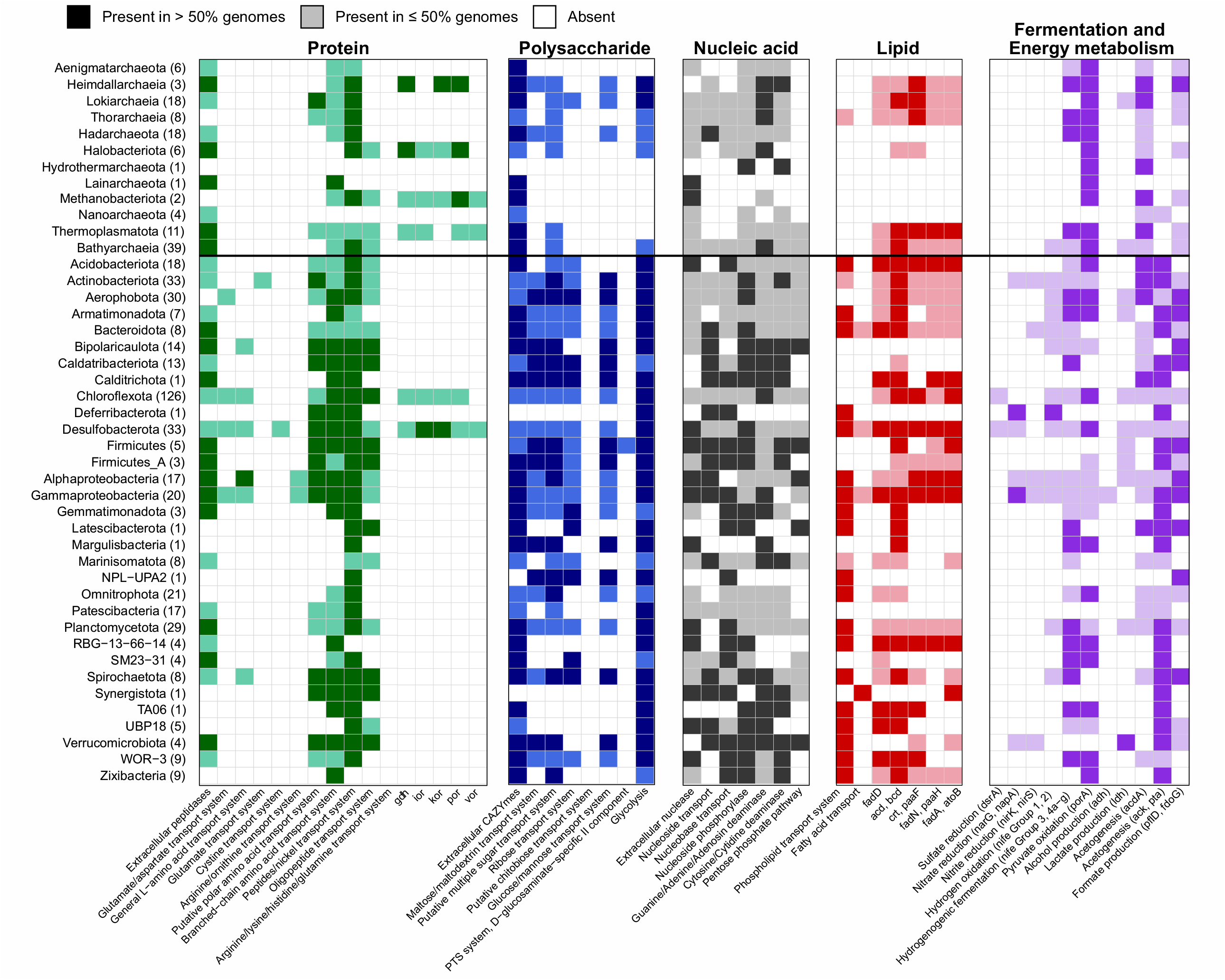
Presence of key genes in degradation of biological macromolecules, fermentation and energy metabolisms detected across phylogenetic clusters. Shading of dark and light colors indicate gene presence in >50% and 1–50% of the MAGs in each phylogenetic cluster, respectively. The number of MAGs per phylogenetic cluster is shown in brackets. The presence of a pathway was determined by the software METABOLIC. A complete list of the annotation is reported in **Table S7**.

**Figure 4.**
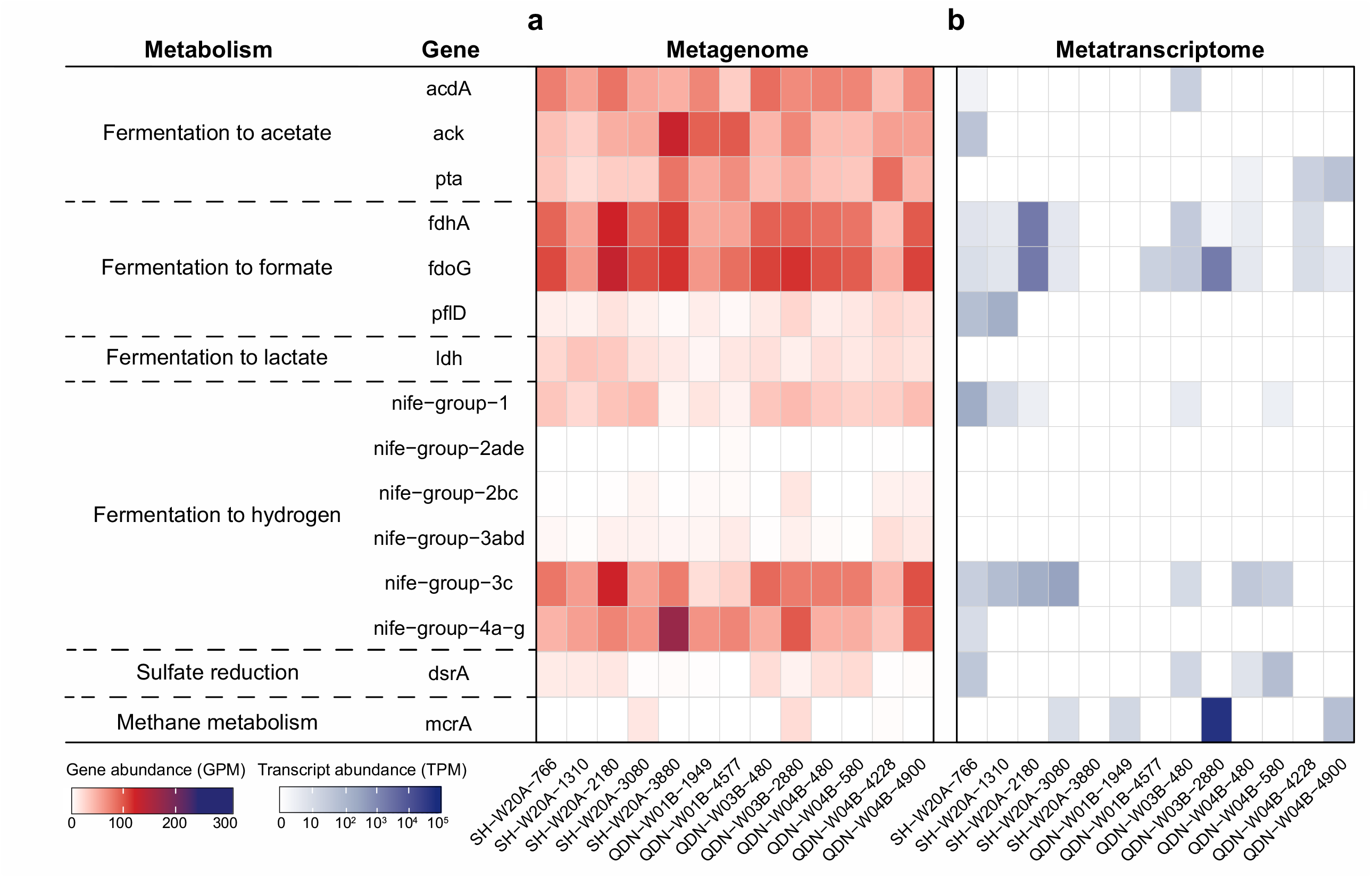
Abundance and expression of metabolic marker genes in the deep subsurface sediments from the gas hydrate zone. (a) The abundance of each gene in metagenomes. (b) The abundance of each transcript in metatranscriptomes. Gene and transcript abundances are represented in the units of genes per million (GPM) and transcripts per million (TPM), respectively.

A majority of the MAGs encodes extracellular peptidases and amino acids transporters (**Figure 3 and Figure S8**). However, only a few lineages such as Chloroflexota, Desulfobacterota and *Hemidallarchaeia* were capable of further metabolizing amino acids (*gdh, por, vor, kor* and *ior*; **Figure 3**), unitizing environmental proteins as energy source. In contrast, most of the subseafloor microbes lacked downstream pathways for amino acid degradation. The amino acids may then be channeled into the salvage pathway used for incorporation into newly synthesized peptides/proteins. Potential DNA-degraders were as diverse as carbohydrate-degraders, including Chloroflexota, Desulfobacterota, Proteobacteria and *Bathyarchaeia*. (**Figure 3)**. They contained genes for extracellular nucleases, DNA polymer breakdown, nucleoside/nuclebase transporters and phosphohydrolases for cleaving nucleoside into the ribose and base moieties, along with pathways for purine and pyrimidine degradation. The released ribose could be further utilized via the pentose phosphate pathway (**Figure 3)**. The retrieved MAGs of Caldatribacteriota and Verrucomicrobiota did not encode genes for extracellular nucleases, but genes for nucleoside/nuclebase transporters and corresponding downstream pathways. Genes coding for lipid degradation were less abundant in the sediments. Chloroflexota, Desulfobacterota, Proteobacteria, Bacteroidota and Actinobacteriota were found to harbor catabolic potentials for lipid degradation (**Figure 3)**. They encoded phospholipid/fatty acid transporters and complete beta-oxidation pathways. The dominant lineage of Caldatribacteriota in the deeper subseafloor seems to lack most genes that encode mechanisms to import and metabolize lipids or fatty acids. Caldatribacteriota MAGs recovered from other anaerobic environments, e.g. petroleum reservoirs [35] and hot spring sediments [36], have also been shown to lack genetic potential for fatty acid degradation.

### Sulfate reducers and methanogens are rare

Only a few MAGs were identified as potential sulfate reducers. They belonged to Chloroflexota (*n* = 5) and Desulfobacterota (*n* = 10, **Figure 3**). These sulfate reducers possess genes encoding a near complete sulfate-reducing pathway (i.e., *dsrA*/*B, aprA*/*B* and *sat*) and other key genes including *dsrC* and *dsrMKJOP* (**Figure S9**). With the availability of sulfate in SMI-above samples, the products of microbial fermentation such as acetate, hydrogen or C1-compounds can be further oxidized by sulfate reducers to CO_2_. A quantitative assessment of *dsrA* genes suggested that potential sulfate reducers are solely detected in sediments above the SMIs (up to 27.12 GPM, **Figure 4a**).

The *mcrA* genes (**Figure 4a**) were only detected in or below the SMIs from QDN-W04B (up to 2.58 GPM), SH-W20A (up to 18.35 GPM), QDN-W03B (up to 25.79 GPM) and QDN-W01B (0.23 GPM). A total of seven recovered MAGs possessed genes encoding McrA. Taxonomic placements showed that these MAGs belonged to *Heimdallarchaeia, Methanomassiliicoccales* and *Methanofastidiosales*, along with ANME-1 and ANME-2 (**Figure S10**). With the exception of *Heimdallarchaeia* SCS_cbin_173, each identified archaeal MAG encoded a complete MCR complex (McrABG) located on a long contig (>10 kbp) with nearby genes being annotated as methane metabolism-related enzymes, as well as tRNAs and ribosomal proteins that were interspersed within the corresponding contig (**Figure S11**). Such an organization of genes was also previously reported in other Mcr-containing archaea [37]. *Heimdallarchaeia* SCS_cbin_173 only encoded partial Mcr complex (McrAG) located on a short contig (3 kbp) without nearby genes mentioned above (**Figure S11**). A low completeness (55.4%) of *Heimdallarchaeia* SCS_cbin_173 is likely responsible for the absence of these genes.

Based on the McrA phylogeny, Mcr complexes in this study were categorized into two major groups (**Figure 5a**), including (1) conventional Mcr clustering with known methanogens or ANME and (2) Mcr-like enzymes associated with anaerobic oxidation of multi-carbon alkanes. *Methanofastidiosales* QDN-W01B-1949_sbin_13 and *Methanomassiliicoccales* QDN-W01B-4577_sbin_7 were closely related to characterized H_2_-dependent methylotrophic methanogens [38, 39] based on both phylogenomic and McrA trees (**Figure 5a and Figure S10**). These two near-complete MAGs (90.26-96.46%) lacked genes encoding for the methyl-H_4_MPT:coenzyme M methyltransferase (Mtr) and methyl-branch of the Wood–Ljungdahl pathway (**Figure S12**), that were present in all CO_2_-reducing methanogens [40]. Instead, they encoded methyltransferases (**Figure S12**) with the potential to support methanogenesis from methanol (MtaA) and methylamine (MtbA). These two hydrogen-dependent methylotrophic methanogens MAGs were found to be present at 19.49 and 45.77 mbsf in QDN-W01B with low abundances (0.09-0.11% of the communities; **Figure S13 and Table S4**).

**Figure 5.**
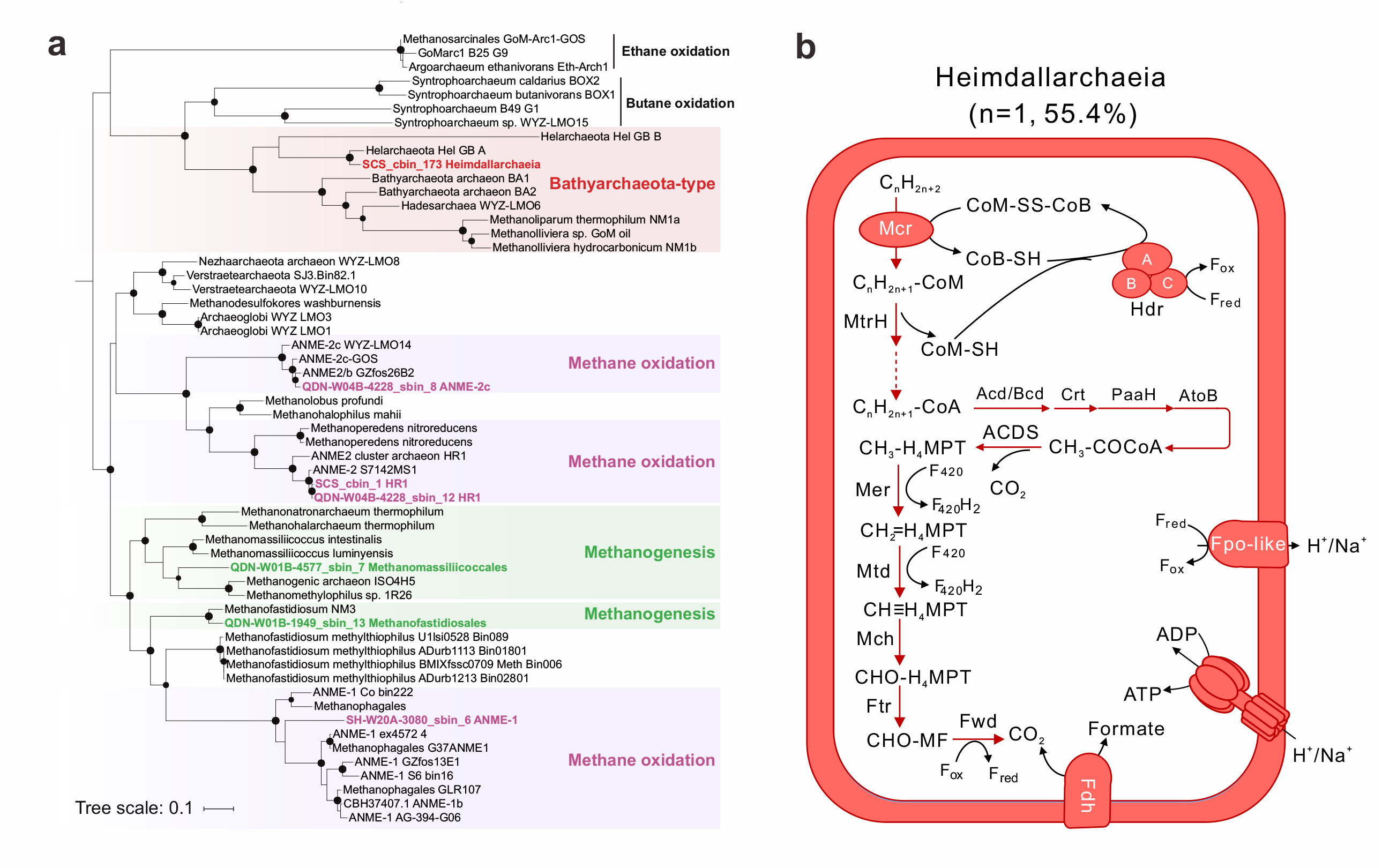
Methane and alkane metabolism. (a) Phylogenetic tree constructed based on alignments of McrA protein sequences. Black dots indicate bootstrap values of 70– 100%. Scale bars indicate the average number of substitutions per site. (b) Anaerobic oxidation of multi-carbon alkanes. Dashed arrows indicate steps catalyzed by unconfirmed enzymes. The percentages between brackets indicate the estimated completeness of the corresponding MAGs. A complete list of metabolic information can be found in Table S11.

Overall, both gene- and genome-resolved approaches demonstrated that the sulfate reducers and methanogens were found to be not abundant in deep subseafloor sediments experiencing methane fluxes.

### Anaerobic methanotrophs and alkanotrophs also are minor members

Four MAGs affiliated with anaerobic methanotrophs ANME-1 and ANME-2 possessed conventional *mcr*, along with an archaeal Wood–Ljungdahl pathway for oxidizing methane to CO_2_, and HdrABC to recycle the CoM and CoB heterodisulfides (**Figure 5a and Figure S14**). ANME-1 was found to be abundant in sulfate-depleted layers, i.e., at 30.8 mbsf in SH-W20A and 28.8 mbsf in QDN-W03B with relative abundances ranging from 2.7% to 3.7% (**Figure S13**). ANME-2 were also found in sulfate-depleted layers i.e., at 42.28 and 49.00 mbsf in core QDN-W04B, but with relatively lower abundance compared to ANME-1 (0.2-1.3%; **Figure S13**). None of ANME MAGs encoded any canonical terminal reductases (such as sulfate and nitrate reductases; **Table S9**), leading to the conclusion that syntrophic partners would be necessary to enable their growth on methane. However, no obvious syntrophic sulfate reducers were identified in these sulfate-depleted sediments (**Figure S13**). All ANME MAGs contained genes for multi-heme cytochromes (**Table S10**) that are often involved in direct electron transfer to a bacterial partner or iron/manganese oxides [44]. Therefore, ANME residing in SMI-below sediments from SH-W20A, QDN-W03B and QDN-W04B might use multi-heme cytochromes to deliver electrons to extracellular metal oxides or metal-reducing bacteria.

One *Heimdallarchaeia* MAG contained *mcrA* sequences (**Figure 5a**) that were clustered with homologs from recently discovered alkane-oxidizing *Helarchaeales* (namely Helarchaeota in NCBI taxonomy) [45, 46]. The McrA from these two lineages formed a monophyletic clade (**Figure 5a**) with those from *Syntropharchaeum* [41], *Bathyarchaeia* (namely Bathyarchaeota) [47], Hadarchaeota (namely Hadesarchaea) [23], and *Ethanoperedens ethanivorans* (namely *Argoarchaeum ethanivorans*) [48], which were all proposed or experimentally proven to perform anaerobic alkane degradation. This MAG also encoded HdrABC for reoxidizing cofactors, methyltransferases to convert alkyl-CoM to acyl-CoA, a complete beta-oxidation pathway to oxidize acyl-CoA to acetyl-CoA, and an archaeal Wood-Ljungdahl pathway (**Figure 5b**), similar to genomic features found in *Helarchaeales* [45, 46]. The *Heimdallarchaeia* MAG present here contained genes encoding Fpo complex for energy transfer across the cell membrane (**Figure 5b**). The identified *Heimdallarchaeia* was rare among the four cores, comprising only 0.002-0.03% of the community (**Figure S13**). Like its *Helarchaeales* relatives, this *Heimdallarchaeia* MAG lacked internal electron sinks and multi-heme cytochromes, but contained formate dehydrogenases that could facilitate the transfer of reducing equivalents in the form of formate (**Figure 5b)**. Genes encoding hydrogenases were not identified in the genome of the *Heimdallarchaeia* (**Table S9**), possibly related to its low genome completeness.

### Gene transcription related to macromolecule utilization and alkanes oxidation

Metatranscriptomic sequencing was performed on these samples to elucidate microbial activities for recycling of macromolecules and alkanes in response to distinct methane fluxes. Genes encoding various extracellular CAZymes were transcribed across most samples. The transcripts encode the breakdown of pectin, chitin, amorphous cellulose and polysaccharides (**Figure 6**). These transcripts for extracellular CAZymes were expressed in Chloroflexota (up to 17 TPM), Caldatribacteriota (up to 2122 TPM), Verrucomicrobiota (up to 8411 TPM), Bipolaricaulota (up to 942 TPM), Planctomycetota (up to 147 TPM), *Bathyarchaeia* (up to 96 TPM), Thermoplasmatota (up to 88 TPM) and Hadarchaeota (up to 1113 TPM). In most samples, many archaea and bacteria transcribe genes encoding various extracellular peptidases, including Chloroflexota, Caldatribacteriota, Bipolaricaulota, Planctomycetota, *Bathyarchaeia*, Thermoplasmatota, *Lokiarchaeia* and Hadarchaeota (**Figure 6**). Although very few transcripts of secretory CAZymes and peptidase were detected in QDN-W01B, we observed transcripts related to various CAZymes and peptidase located in the cell membranes or cell walls (cell-attached CAZymes and peptidase) in this core (**Table S12**). These cell-attached CAZymes and peptidases are also important for hydroxylation of environmental macromolecules via a tighter hydrolysis-uptake coupling, especially in the deep subseafloor environment where organic matter is refractory [34]. Both gene- and genome-resolved analyses revealed that genes involved in sulfate reduction (*dsrA*) were only transcribed in sediments above SMI, with Desulfobacterota and Chloroflexota being major sulfate reducers (**Figures 4 and 6**). This result also illustrated that Chloroflexota and Desulfobacterota in SMI-above sediments (**Figure 2**) most likely actively degraded macromolecule-derived carbon coupled with sulfate reduction. When sulfate was depleted, transcripts assigned to formate, acetate, and hydrogen fermentation were found to be highly abundant, e.g. in SH-W20A and QDN-W03B (**Figure 4b**).

**Figure 6.**
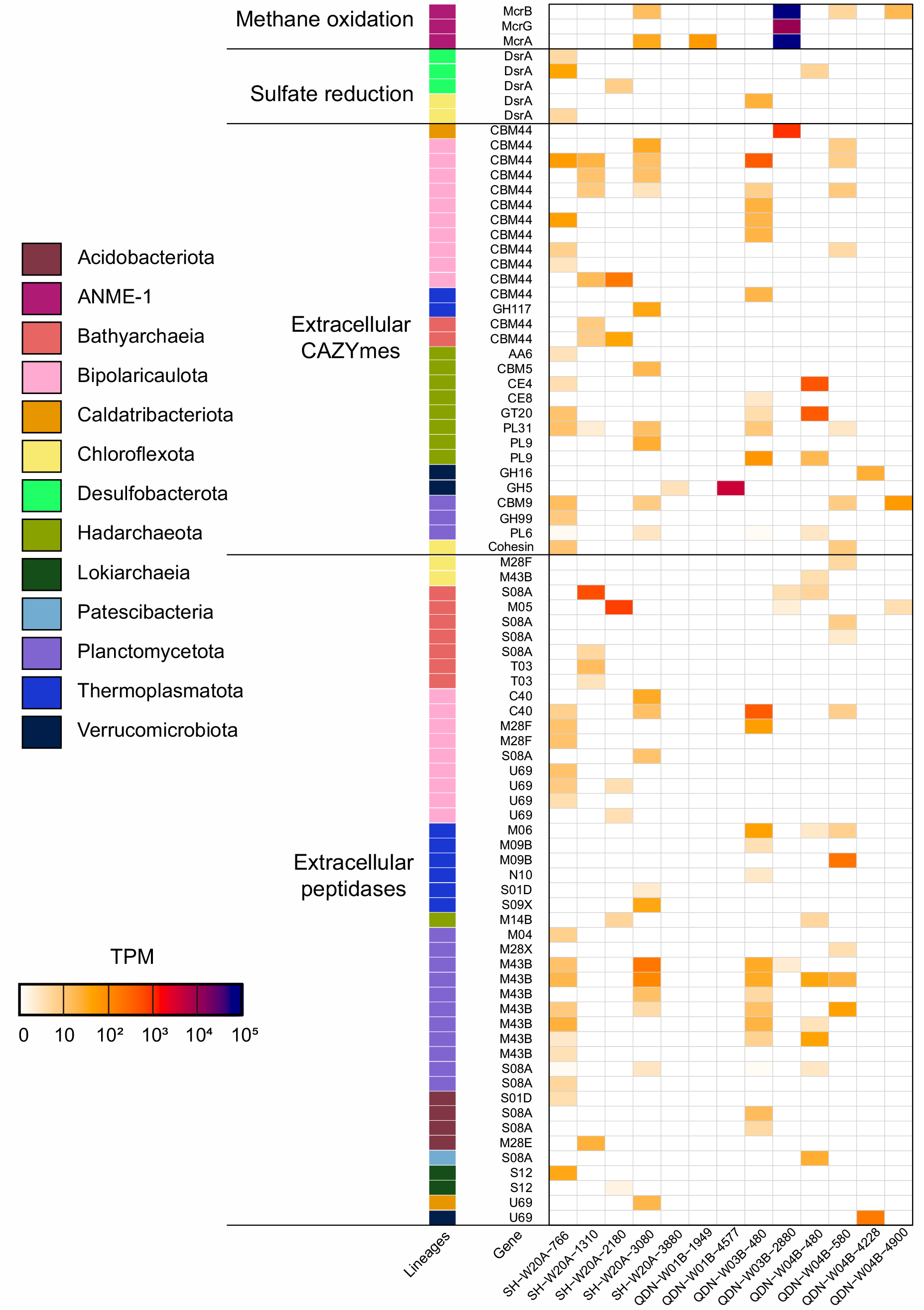
Transcription of genes for sulfate reduction, methane oxidation as well as extracellular CAZymes and peptidases in the genomic bins. The expression levels of each gene were represented in the unit of transcripts per million (TPM). A complete list of the transcriptional information is reported in **Table S12**.

Genes coding *mcrA* gene were expressed in and below the SMI zones (**Figure 4b**), with highest transcription in QDN-W03B at 28.8 mbsf (71981 transcripts per million, TPM). Most of the *mcrA* transcripts mapped the MAG ANME-1 SH-W20A-3080_sbin_6 (**Figure 6 and Table S12**) in QDN-W03B (139474 TPM at 28.8 mbsf), whereas this transcription was much lower in SH-W20A (81 TPM at 30.8 mbsf). Genes encoding beta (McrB) and gamma (McrG) subunits of Mcr complex encoded in this genome were also expressed at different degrees. Notably, *mcrA* genes were hardly detected in the metagenomes and MAGs of QDN-W01B and the transcriptome contained only few *mcrA* reads. These results confirmed the occurrence of ANME archaea in SMI as deep as ∼30 mbsf, whereas methane metabolism is very rare in sediments above SMIs.

## Discussion

This study combined geochemistry, metagenomics and metatranscriptomics to characterize deep subseafloor microbial community structures and carbon metabolism in sediments experiencing methane fluxes. Methane fluxes strongly imprint the geochemistry of porewaters along depths in the Shenhu area and Qiongdongnan Basin. A shallow depth of SMI was observed in QDN-W01B corresponding to a rapid sulfate depletion, which may be attributed by high methane fluxes [6]. In SH-W20A, QDN-W03B and QDN-W04B, low methane fluxes allowed sulfate diffuseinto greater depth.

Metagenomic profiling revealed that microbial community compositions in deep subseafloor sediments significantly correlated with the redox zonation. In SMI-above subseafloor layers, members of Chloroflexota predominated, while members of Caldatribacteriota were dominant in SMI-below subsurface sediments. Chloroflexota and Caldatribacteriota are broadly defined taxonomic groups and have been found to co-occur at high abundances in sediments from many different regions across global oceans [49]. The transition of Chloroflexota to Caldatribacteriota between redox zones along with sediment depth implies intense subseafloor selection [50]. Members of Caldatribacteriota are likely more adaptive to deep anoxic sediments with less organic carbon and energy availability [3, 51], which explains their successful survival in the deep subseafloor.

The genome-based functional analysis suggests that microbial communities utilize biologically produced macromolecules (mainly carbohydrates) as carbon and energy sources regardless of the availability of methane. The main heterotrophs are Chloroflexota, Caldatribacteriota, Verrucomicrobiota, Hadarchaeota and Bathyarchaeia. Our results are similar to what were observed in common marine sediment not influenced by methane fluxes. For example, metatranscriptomics and enzyme assays suggested that Caldatribacteriota actively catabolized sugars and protein in the subsurface sediments of Baltic Sea [52]. Verrucomicrobiota and Chloroflexota were also found to be dominant heterotrophs in sediments of the Helgoland mud area [14]. Most of these microorganisms seem to be generalists that degrade a wide range of biopolymers. The majority of the deep subseafloor communities could extracellularly hydrolyze macromolecules by secreting enzymes into ambient environments and take up oligomers via various transporters. Genes encoding for CBM44 involved in cellulose and xyloglucan binding [53] were widely transcribed in the deep subseafloor communities. Cellulose and xyloglucan are derived from terrestrial vascular plants [54]. In short, our results demonstrate the degradation of plant matter into the deep subseafloor sediments from the gas hydrate zone.

Nucleic acids represent another key trophic resource and provide an energy source [55], but their role in sedimentary biogeochemical cycles has largely been ignored to date. The catabolism of nucleic acids involved in extracellular cleavage and transmembrane import before intracellular degradation [17, 18]. Many of the Caldatribacteriota and Verrucomicrobiota contain nucleoside/nuclebase transporters for the uptake of these compounds, but they lack extracellular nucleases such as Nuc superfamily (DNA/RNA nonspecific endonucleases) [56]. These microorganisms act as opportunists which depend on other heterotrophs that produce exoenzymes. Such a co-occurrence of exoenzyme-producing and opportunistic scavengers seem to be common in nature and have for instance observed in Guaymas Basin hydrothermal sediments [17] and terrestrial soils [57].

The products of fermentation (formate, lactate, acetate and hydrogen etc) can be further oxidized into CO_2_ by other microbes using sulfate as electron acceptor in the SMI-above layers or used for methanogenesis when sulfate was depleted. Except for the typical sulfate reducers of Desulfobacterota [58], the presence of *dsrAB* genes in several MAGs (*n* = 5) of Chloroflexota suggested their ability for sulfate respiration. Bacteria of Chloroflexota are widespread in various marine sediments with high abundance [59], which display a broad spectrum of metabolic traits, e.g. reductive dehalogenation and fermentation of organic matter [60, 61]. Only recent year has their roles in the deep subsurface sulfur cycle been recognized [62]. Our data expand the limited number of Chloroflexota genomes, which may provide important information for the cultivation of deep-sea sulfate-reducing Chloroflexota. Although the condition in the subseafloor was considered to be suitable for hydrogenotrophic or acetoclastic methanogenesis, we only detected H_2_-dependent methylotrophic methanogens (*Methanomassiliicoccales* and *Methanofastidiosales*), which produce methane using hydrogen, C1 substrates and synthesize biomass using acetate [63]. Similarly, in other reports about the sediments of JiaoLong, Shenhu and Haima methane seep from the South China Sea as well as hydrate-bearing sediments from eastern Nankai Trough, the H_2_-dependent methylotrophic methanogenesis was found to be a major methanogenic pathway [64-67]. The methylotrophic methanogenesis were reported to be able to unitize non-competitive substrates (methanol, methylamines, or methyl sulfide and so on) originated from the degradation of organic macromolecules such as lignin and pectin [68]. Therefore, the fermentation of macromolecules does not necessarily support hydrogenotrophic or acetoclastic methanogens but can also be linked to H_2_-dependent methylotrophic methanogenesis.

The upward methane fluxes serve as alternative carbon and energy source for a small part of ANME community. Combined metagenomic and metatranscriptomic evidence suggested that higher gene and transcript abundances for *mcrA* genes were found in the SMI-below sediments of SH-W20A, QDN-W03B and QDN-W04B. ANME-1 (up to 3.7% of the whole community) and ANME-2 (up to 1.3%) were found to be the main methane oxidizers in SMI-below sediments, with ANME-1 being highly active based on expressed transcripts. The observed response of ANME communities in the deep subseafloor is different from those fueled by high methane fluxes in the sulfate-rich surface sediments from Arctic seafloor gas hydrate mounds [6]. Previous observations of ANME-1 and ANME-2 clades have focused mainly on shallow sediments at no more than 20 mbsf, in the sulfate-methane transition zones (SMTZ) or occasionally sulfate-depleted horizons below SMTZ [6, 69-72]. The results presented here indicate that ANME archaea are still active in deeply buried marine sediments with depths over 30 mbsf. Despite in sulfate-depleted subseafloor layers, the presence of genes for multi-haem cytochromes in these ANME archaea points to that they might use solid metal oxides (e.g. Fe and Mn) other than sulfate for energy conservation [73], coupled with oxidizing methane migrated from below reservoirs through gas chimneys.

One *Heimdallarchaeia* MAG stood out particularly because it contained alkyl-CoM reductase and downstream parts of the utilization of short-chain alkanes. Despite belonging to the rare biosphere (up to 0.03% of the community), *Heimdallarchaeia* may play a substantial role in the community in response to environmental oxidation of non-methane alkanes. Mcr-based methane cycling was thought to be limited to Halobacteriota (namely Euryarchaeota) [74]. Until recently, non-Halobacteriota phyla, including TACK superphylum and *Helarchaeales*, have been shown to contain proteins with homology to Mcr [42, 45, 47]. To the best of our knowledge, *Helarchaeales* is so far the only known group of Asgard archaea genetically capable of Mcr-mediated hydrocarbon oxidation. Based on the phylogenetic analysis, *mcrA* genes from *Heimdallarchaeia* and *Helarchaeales* are closely related (Figure 5). Furthermore, both *Heimdallarchaeia* and *Helarchaeales* lack genes for canonical pathways of sulfate reduction and cytochromes. They hence most likely need external partners to consume the reducing equivalents in the forms of hydrogen and formate [45, 46]. The *mcr* genes in *Helarchaeales* were previously thought to acquire from e.g. *Bathyarchaeia* or *Syntrophoarchaeum* due to horizontal gene transfer events. Within the Asgard archaea, these genes are restricted to *Helarchaeales* [45]. The discovery of *Heimdallarchaeia* with potential for multi-carbon alkane degradation indicates a broader view of the evolution and expansion of hydrocarbon oxidation pathways within Asgard archaea [45, 46, 75, 76].

Together, our results uncover metagenomic blueprints and metatranscriptomic activities of microbes residing in deep subseafloor sediments from the gas hydrate zone. The sulfate above the SMI sustains small numbers of sulfate reducers. The upward methane does not always provide energy and carbon source for microbial communities due to lack of suitable electron donors in gas hydrate sediments. More importantly, these findings underpin that fermentation of macromolecules sustains a large number of different archaea and bacteria in deep subseafloor sediments. Microorganisms that use the fermentation products for methanogenesis or sulfate reduction, are of minor abundance and their marker genes such as *mcr* or *dsr* are in low expression. This shows that energy conservation is concentrated in the fermentative organisms. Albeit the fermentation reaction cannot be directly recognized by in porewater profiles, in the studied sediments the fermentative microorganisms outnumber the sulfate reducers and methanogenesis by an estimated factor of the combination of organic matter, sulfate and upward methane.

## Material and methods

### Sampling sites

The studied sediments were drilled from the Qiongdongnan Basin and Shenhu area (**Figure S1**), where considerable amounts of gas hydrates were discovered [20, 77, 78]. The Qiongdongnan Basin is an oil-bearing, fault-depression structural basin on the northwestern continental shelf of the South China Sea [20]. The Shenhu area is in the middle of the northern slope of the South China Sea, tectonically located in Pearl River Mouth Basin [77]. Four cores penetrating to 100−188 mbsf were analyzed in this study, with water depths ranging from 1000 to 1500 meters. They were obtained during the 2019 gas hydrate drilling expedition (GMGS6) conducted by Guangzhou Marine Geological Survey. Detailed descriptions on seismic data, cores, and grain sizes of these samples were published previously [26]. Three drilling sites of cores QDN-W01B, QDN-W03B and QDN-W04B were located in the Qiongdongnan Basin, and the other core SH-W20A was recovered from the Shenhu area. Each sediment core was sectioned using a core extruder on board immediately following their retrieval. Samples were then frozen at −80 °C for subsequent geochemical analysis and nucleic acid extraction.

### Geochemical measurements

Porewater geochemistry and total organic matter content were analyzed for 67 samples at the depth of approximately 1 to 79 mbsf, and 45 samples ranging from 0.5 to 73.6 mbsf were selected for the headspace gas analysis, following methods reported in our previous studies [75, 79]. Briefly, methane cconcentrations were measured using a headspace equilibration technique by an Inficon Fusion MicroGC gas chromatograph with molecular sieve and PLOT Q columns and thermal conductivity detectors. Sulfate concentrations were determined using ion chromatography (Metrohm 790 Personal IC). Concentrations and δ^13^C values of porewater DIC were analyzed via a continuous flow mode-isotope ratio mass spectrometry (CF-IRMS). The TOC contents of bulk sediment samples were quantified on an elemental analyzer-isotope ratio mass spectrometer (EA−IRMS) after removal of inorganic carbon.

### DNA extraction and metagenomic sequencing

Genomic DNA was extracted from approximately 10 g of sediments for each depth using PowerMax Soil DNA Isolation Kit (Qiagen) according to the manufacturer’s instructions. DNA concentrations were evaluated using a Qubit 4.0 Fluorometer (Thermo Fisher Scientific). Metagenomic libraries were prepared for 13 samples following the manufacturer’s instructions (Illumina Inc.). Sequencing was performed on an Illumina NovaSeq 6000 platform with a 2 × 150 bp paired-end run at Berry Genomics Co. Ltd., Beijing.

### Metagenomic assembly and binning

Raw reads derived from the 13 metagenome libraries were quality-controlled using Read_qc module (-skip-bmtagger) within the metaWRAP v1.2.2 pipeline [80]. Quality-controlled reads were assembled individually using metaSPAdes v3.13.0 [81] and co-assembled using Megahit v1.1.3 [82] within the metaWRAP Assembly module, producing 14 assemblies. Each assembly was binned using Binning module (parameters: -maxbin2 -concoct -metabat2) and consolidated using Bin_refinement module (parameters: -c 50 -x 10) within the metaWRAP pipeline. All binning results were aggregated and de-replicated using dRep v2.5.4 [83] at 95% and 99% average nucleotide identities, for species and strain levels, respectively. Completeness, contamination, and heterogeneity of MAGs were estimated using CheckM v1.0.18 [84].

### Taxonomic assignments of MAGs

The taxonomy of each MAG was assigned using GTDB-Tk v1.3.0 with reference to GTDB 05-RS95 [85]. The assignments were confirmed by visual inspection of taxonomic trees. Reference genomes accessed from NCBI GenBank and the MAGs from this study were used to construct the phylogenomic tree based on concatenation of 43 conserved single-copy genes extracted by CheckM v1.0.18 [84], following the procedures reported in our previous study [13]. The maximum-likelihood phylogenomic tree was built using RAxML v8 with the PROTCATLG model, bootstrapped with 1000 replicates [86].

### Diversity and community profiling

Alpha and beta diversity of microbial communities were carried out using vegan package v2.5. Operational taxonomic units (OTUs) were extracted from the metagenomic data using SingleM v0.12.1 (https://github.com/wwood/singlem) by aligning to a database of 14 single-copy ribosomal proteins [87]. OTU tables were then summarized via rarefying and clustering using SingleM summarise. Shannon index and Chao1 were calculated from the SingleM OTU tables across each of the 14 single-copy marker genes using vegan package v2.5. For beta diversity analysis, Bray-Curtis dissimilarity generated from the *rplB* OTU table was visualized using non-metric multidimensional scaling (NMDS) plots. A pairwise analysis of similarities (ANOSIM) was used to test for significant differences in community similarity between redox zones.

Microbial community structures were determined using both gene- and genome-centric approaches. For better community profiling in gene-centric analyses, 16S rRNA gene fragments (i.e., 16S miTags) were extracted from metagenomic raw reads using the phyloFlash v3.4 pipeline (parameters: -almosteverything) and classified with the SILVA v138.1 database [88]. For genome-centric analyses, the relative abundance of MAGs depreciated at species level were calculated using CoverM v0.4.0 (https://github.com/wwood/CoverM) with the genome mode (parameters: - min-read-percent-identity 0.95 -min-read-aligned-percent 0.75 -trim-min 0.10 -trim-max 0.90).

### Functional annotations

For all contigs assembled from 13 metagenomic samples, the functional annotation was undertaken with METABOLIC v4.0 [89]. All predicted coding sequences were pooled and clustered at 95% nucleotide sequence similarity using CD-HIT v4.8.1 [90] (parameters: -c 0.95 -T 0 -M 0 -G 0 -aS 0.9 -g 1 -r 1 -d 0). Finally, a total of 3,701,930 non-redundant gene clusters were obtained and used as the reference gene catalog for microbial communities. The program Salmon v1.5.0 [91] in the mapping-based mode (parameters: -validateMappings -meta) was used to calculate gene abundance from the reference gene catalog in different metagenomes. Gene abundances were expressed as genes per million (GPM).

For individual MAGs, metabolic genes were identified by METABOLIC v4.0 [89]. Genomes were also annotated using DRAM with default parameters [92] against KOfam, MEROPS and dbCAN databases to identify CAZymes, peptidases, lipases, nucleases, transporters, and other proteins of interest. For cytochrome C detection, MAGs with identified *mcrA* gene were screened for proteins based on characteristic cytochrome C CXXCH domains following the criteria described elsewhere [70]. Protein localization was determined for CAZymes, peptidases and cytochrome C using the web tool Psortb v3.0.3 [93].

### Phylogenies of functional genes

For each gene, amino acid sequences from the current study were aligned with reference sequences using MAFFT v7.471 [94] (–auto option) and trimmed using TrimAl v1.2.59 [95] (–gappyout option). Maximum likelihood trees were constructed using IQ-TREE v2.0.5 [96], implemented in the CIPRES web server, with best-fit models and 1000 ultrafast bootstrap.

### Metatranscriptomic analysis

Total RNA was extracted from replicate samples of the metagenome analysis using the RNeasy PowerSoil Total RNA kit (Qiagen) according to the manufacturer’s instructions. RNA purity and concentration were evaluated using Qubit (Thermo Fisher Scientific). RNA integrity was accurately detected using the Agilent 4200 system (Agilent Technologies). Whole transcriptome amplification of total RNA was carried out using RNA REPLI-g Cell WGA & WTA Kit (Qiagen) according to its protocol. To enrich messenger RNA (mRNA), ribosomal RNA was depleted from total RNA using ALFA-SEQ rRNA depletion Kit. Whole mRNAseq libraries were generated by Guangdong Magigene Biotechnology Co. Ltd. (Guangzhou, China) using NEB Next Ultra Nondirectional RNA Library Prep Kit for Illumina (New England Biolabs) following manufacturer’s recommendations. The constructed libraries were sequenced on an Illumina Novaseq 6000 platform and 150 bp paired-end reads were generated.

Raw metatranscriptomic reads were quality filtered in the same manner as for metagenomes. The reads corresponding to ribosomal RNAs were removed using SortMeRNA v.4.2.0 [97]. Subsequently, these high-quality metatranscriptomic reads were mapped to the predicted protein-coding genes from all the MAGs and the reference gene catalog using Salmon v.1.5.0 [91] in mapping-based mode (parameters: -validateMappings -meta). The expression level for each gene was normalized to transcript per million (TPM).

## Data availability

All metagenomic and metatranscriptomic raw reads used in this study can be accessed at the Sequence Read Archive under BioProject accession number PRJNA739005. The assemblies, reference gene catalog, all MAGs and phylogenetic trees can be found in figshare (https://figshare.com/account/home#/projects/116454).

## Acknowledgments

We thank Tingting Chen, Ling-Dong Shi and Jiwei Li for helpful discussions, and all of those who contributed to the 2019 gas hydrate drilling expedition of China National Gas Hydrate Program.

## Author contributions

XD, JHW and YXF designed this study. XD and CZ analyzed metagenomic and metatranscriptomic data. CZ and TZ prepared the figures. YXF, HL, ZK and XPX were responsible for sediment sampling and performed biogeochemical measurements. XD, CZ, XY and GW interpreted the data. XD, CZ, XY, GW and YXF wrote the paper, with input from other authors.

## Ethics declarations

### Ethics approval and consent to participate

Not applicable.

### Consent for publication

Not applicable.

