## Supplementary Figures for "The majority of microorganisms in gas hydrate-bearing subseafloor sediments ferment macromolecules"

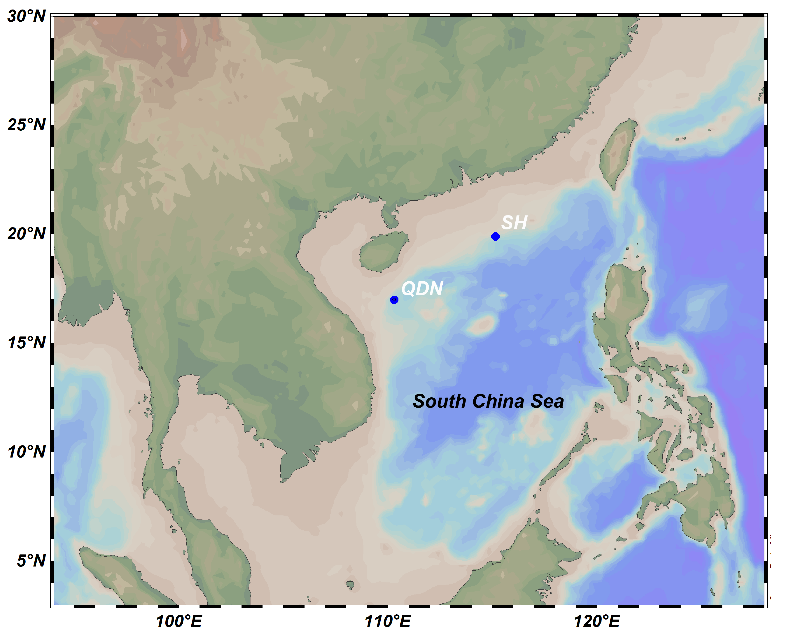

**Figure S1. Location of sampling sites in the gas hydrate zone of Shenhu area (SH) and Qiongdongnan (QDN) Basin from the South China Sea.** All the four drilling sites were located above deep subsurface gas chimneys. They were retrieved from the Shenhu area (n = 1; SH-W20A) and Qiongdongnan Basin (n = 3; QDN-W01B, W03B and W04B). These cores were penetrated to 100−188 mbsf, with water depths ranging from 1000 to 1500 meters.

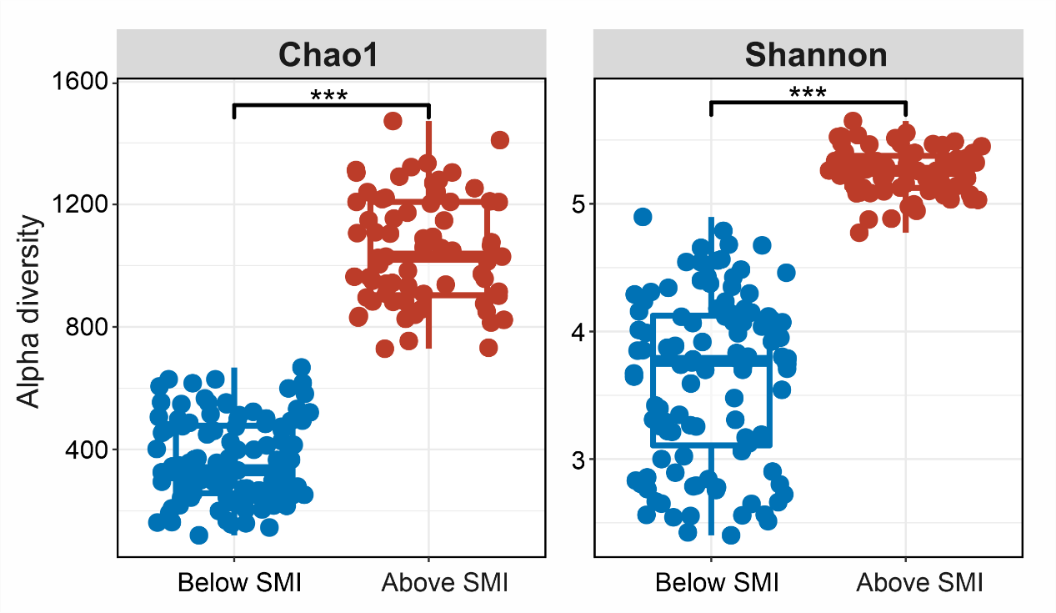

**Figure S2. Comparison of Chao1 and Shannon indices of the microbial community between the SMI-above and -below sediments in the gas hydrate zone.** Chao1 and Shannon indices were calculated based on the SingleM ((https://github.com/wwood/singlem) OTU tables for 14 universal single-copy genes. P-values of differences between redox zones were calculated using Wilcoxon rank sum test. Asterisks denote significance (*** for *P* < 0.001).

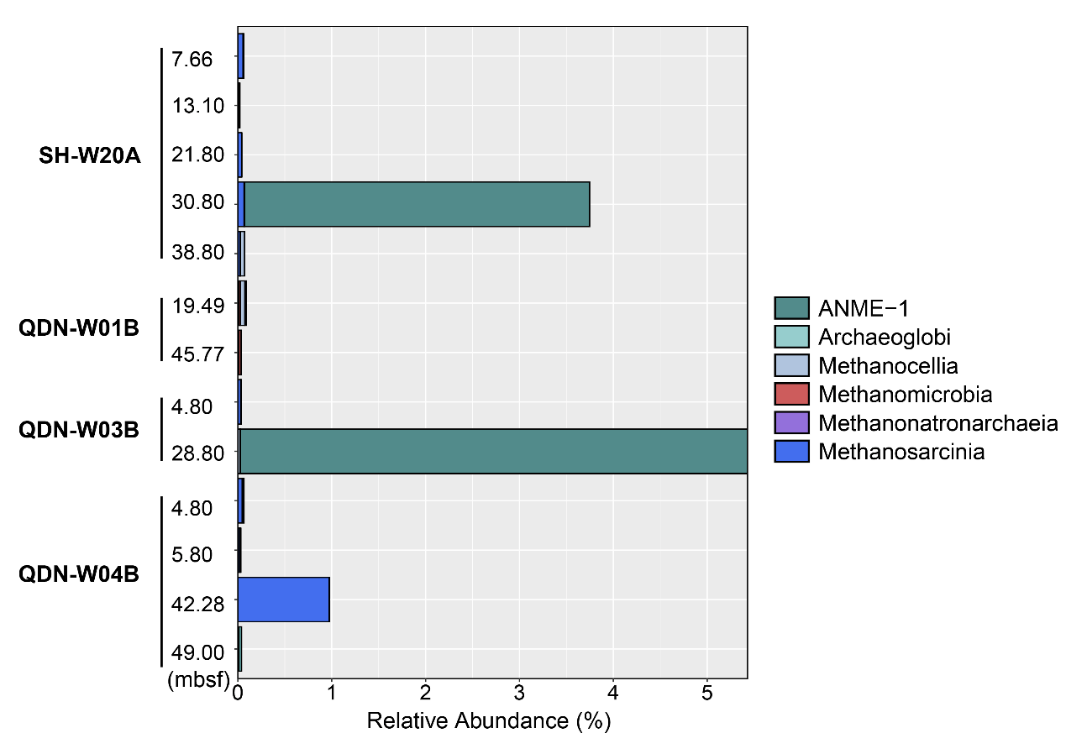

**Figure S3. Relative abundances of each class from the Halobacterota phylum inferred from 16S rRNA gene fragments in metagenomes**.

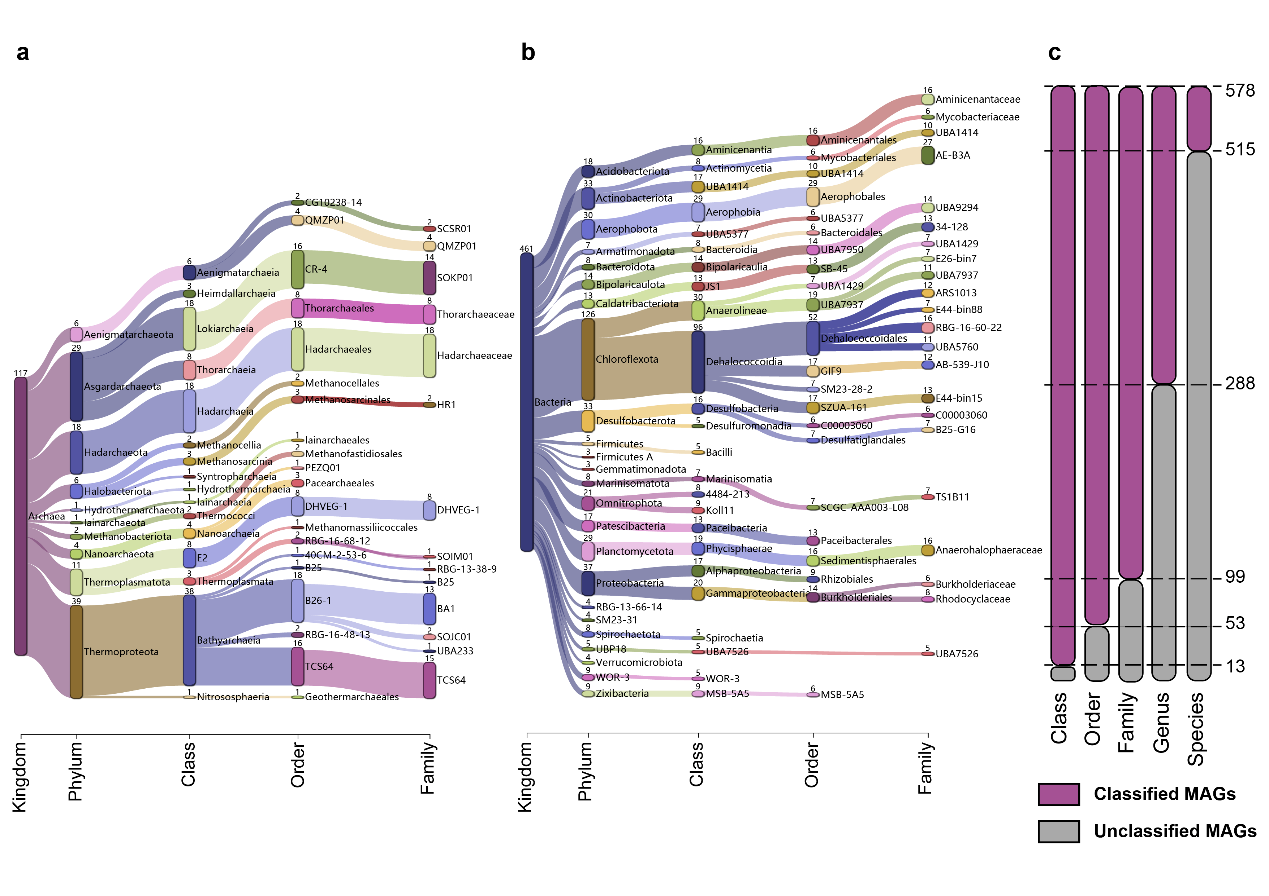

**Figure S4. MAG recovery information across different taxonomic levels**. A Sankey diagram based on assigned GTDB taxonomy showing archaeal (a) and bacterial (b) MAGs at different phylogenetic levels. Numbers indicate the number of MAGs recovered for this lineage. (c) Total MAGs unclassified by GTDB-Tk at each taxonomic level. MAGs were dereplicated at strain level (i.e., 95% ANI). Detailed statistics for 578 MAGs are provided in **Table S3**.

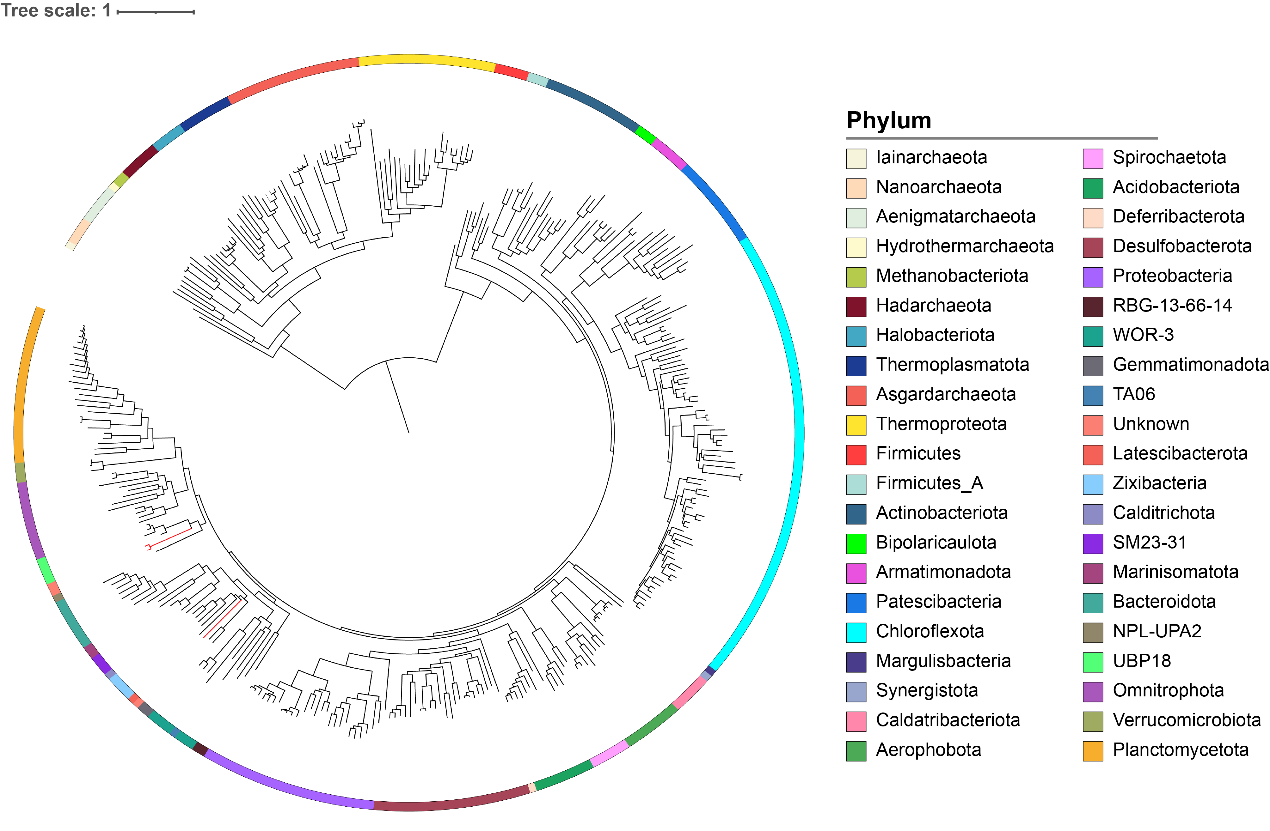

**Figure S5. Phylogenetic placement of 349 MAGs for microbial communities in the deep subsurface sediments from the gas hydrate zone.** Branches in red indicate unclassified MAGs. The maximum-likelihood phylogenomic tree was built based on concatenated amino acid sequences of 43 conserved single-copy genes using RAxML with the PROTCATLG model. The scale bar represents the average number of substitutions per site. Relative abundances of those microorganisms can be found in **Table S4**.

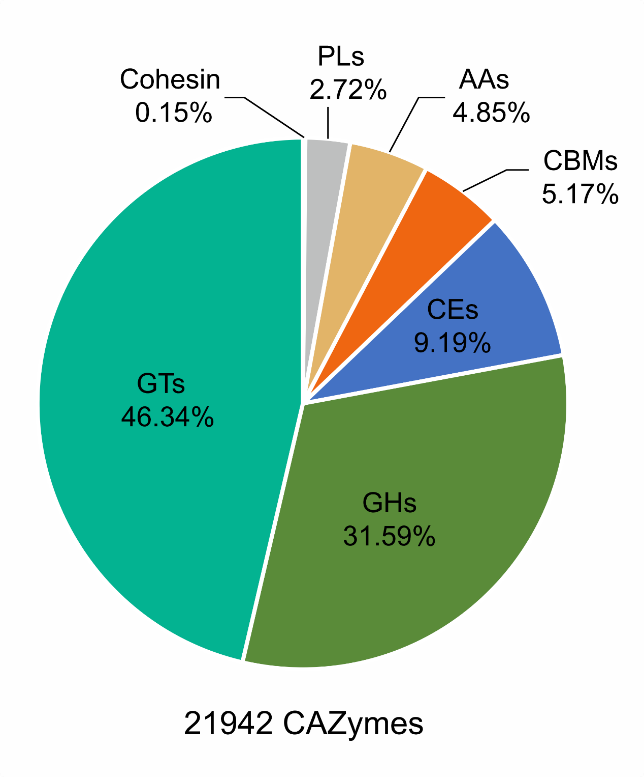

**Figure S6. Carbohydrate-active enzyme composition of the deep subseafloor microbiome from the gas hydrate zone.** GTs: glycosyltransferases; GHs: glycoside hydrolases; CEs: carbohydrate esterases; CMBs: carbohydrate-binding modules; AA: auxiliary activities; PLs: polysaccharide lyases.

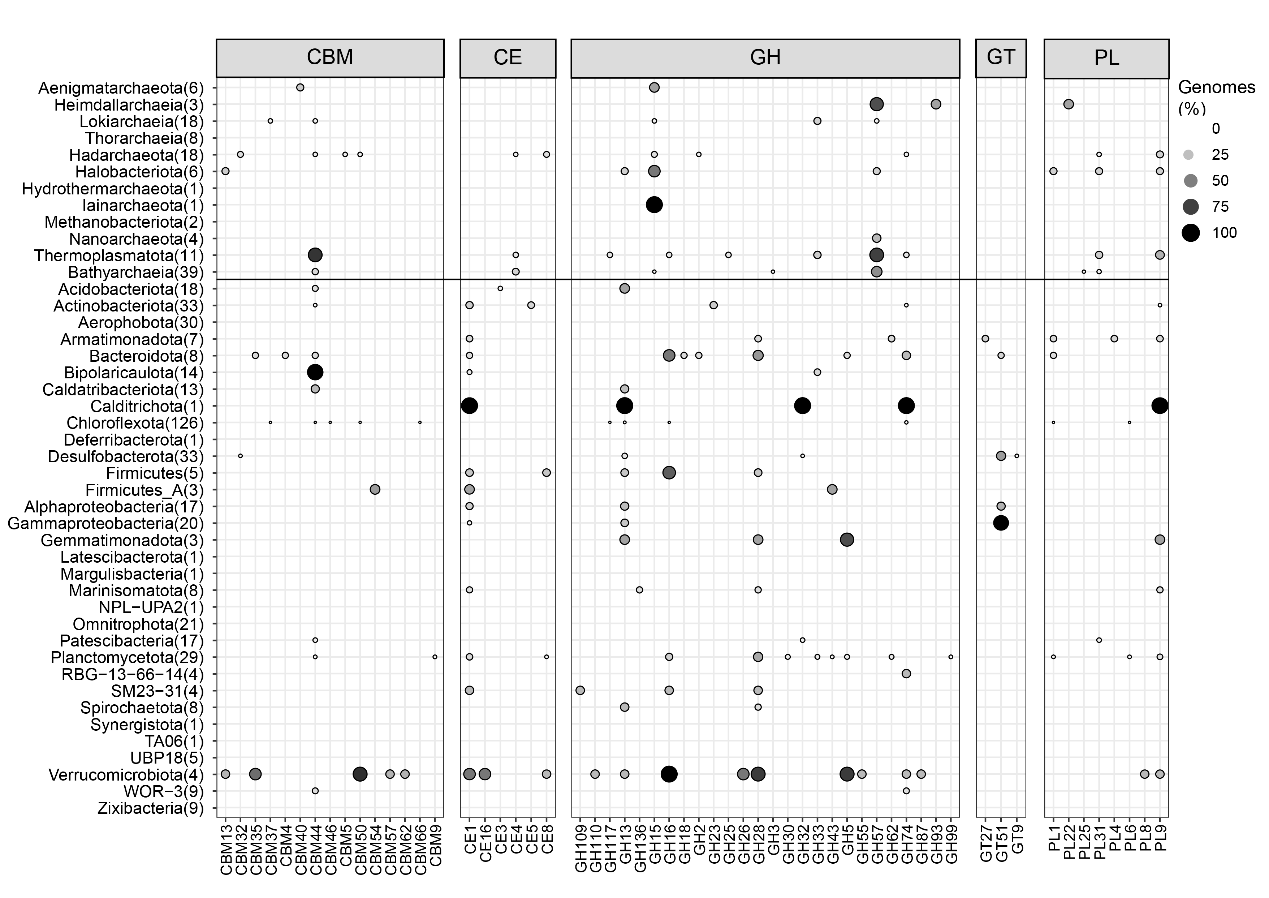
 **Figure S7. Percentage of extracellular carbohydrate-active enzymes (CAZymes) encoded in each phylogenetic cluster.** The number of MAGs per phylogenetic cluster is shown in brackets. CMBs: carbohydrate-binding modules; CEs: carbohydrate esterases; GHs: glycoside hydrolases; GTs: glycosyltransferases; PLs: polysaccharide lyases.

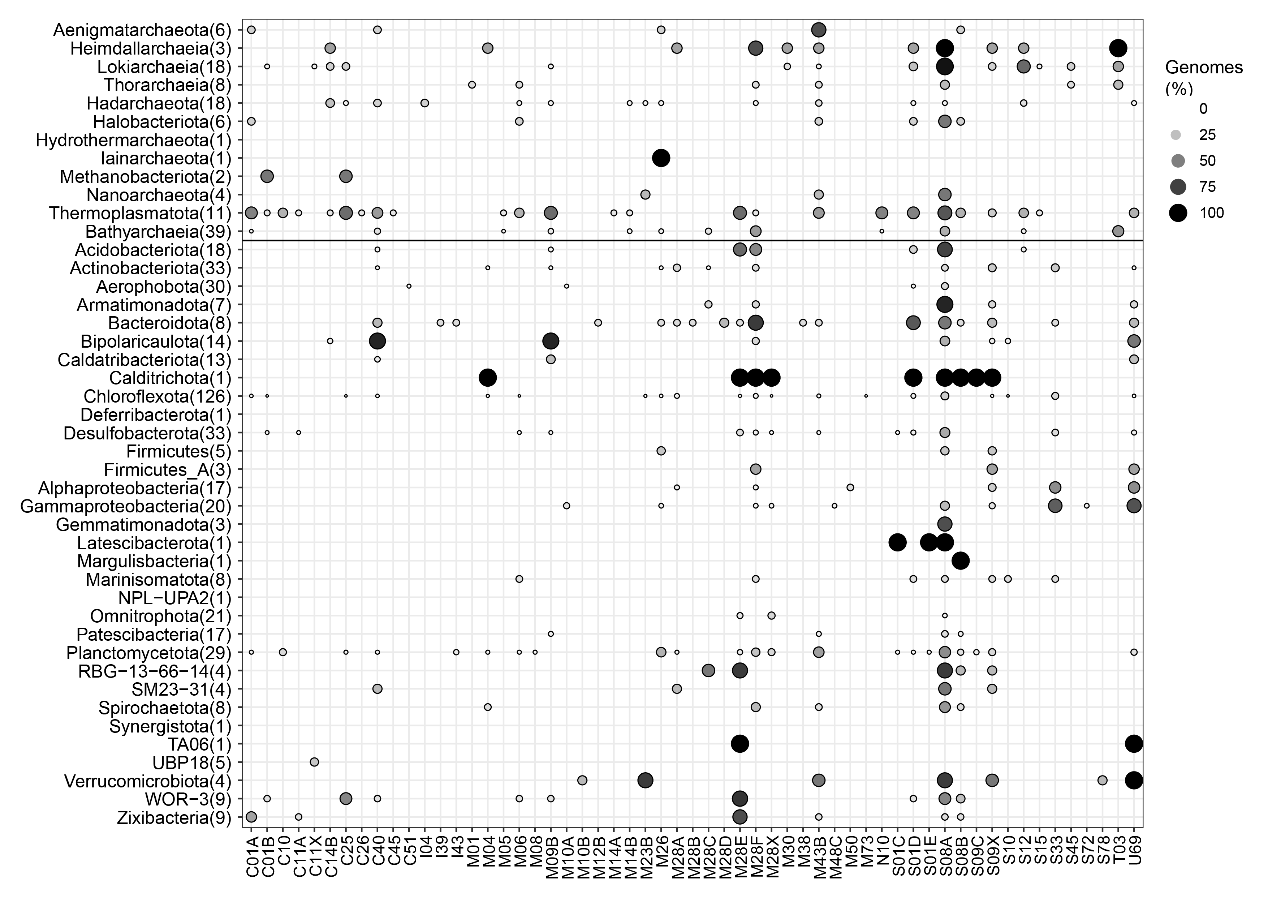

**Figure S8. Percentage of extracellular peptidases encoded in each phylogenetic cluster.** The number of MAGs per phylogenetic cluster is shown in brackets.

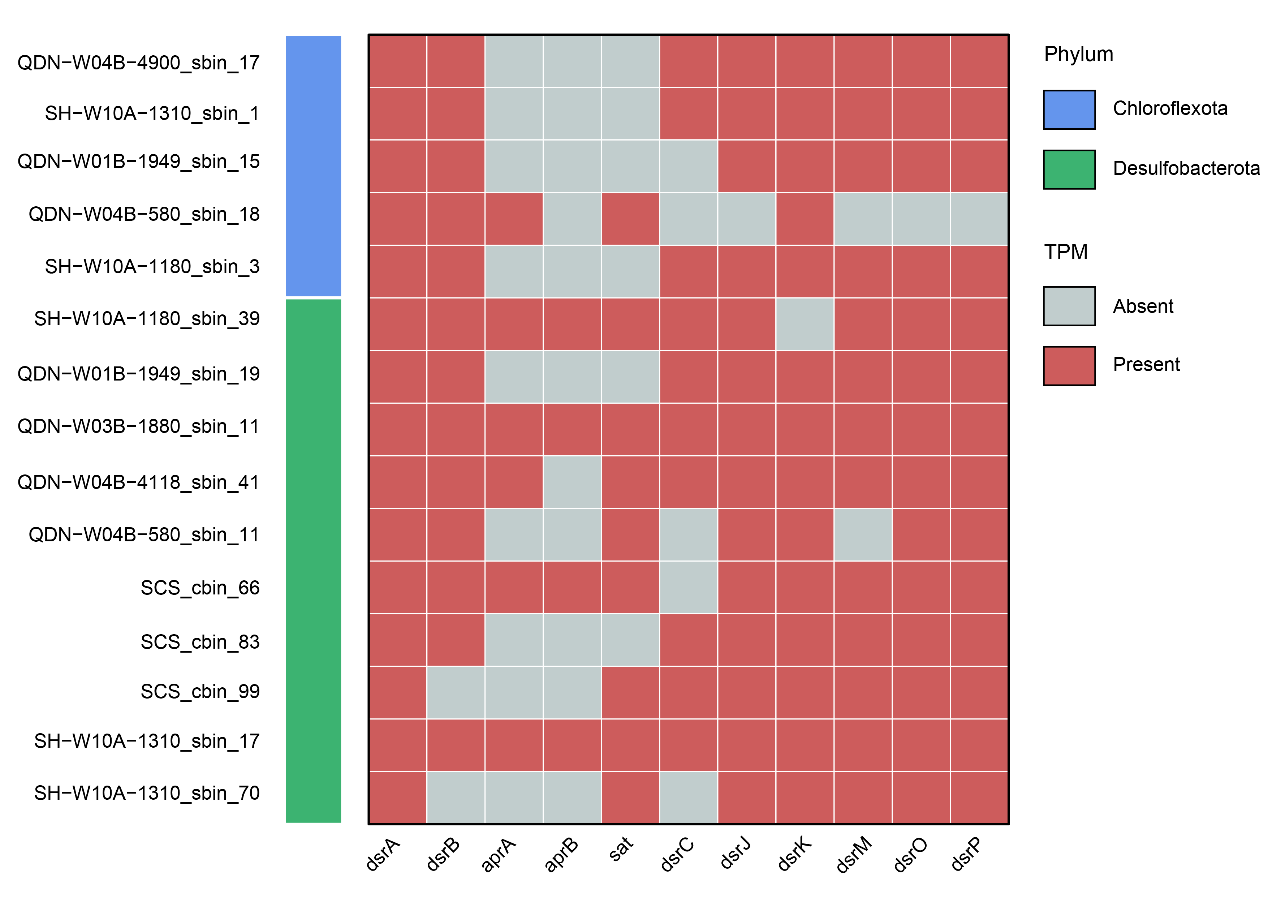

**Figure S9. Presence and absence of genes in sulfate-reducing pathway for recovered sulfate reducers reported in this study.**
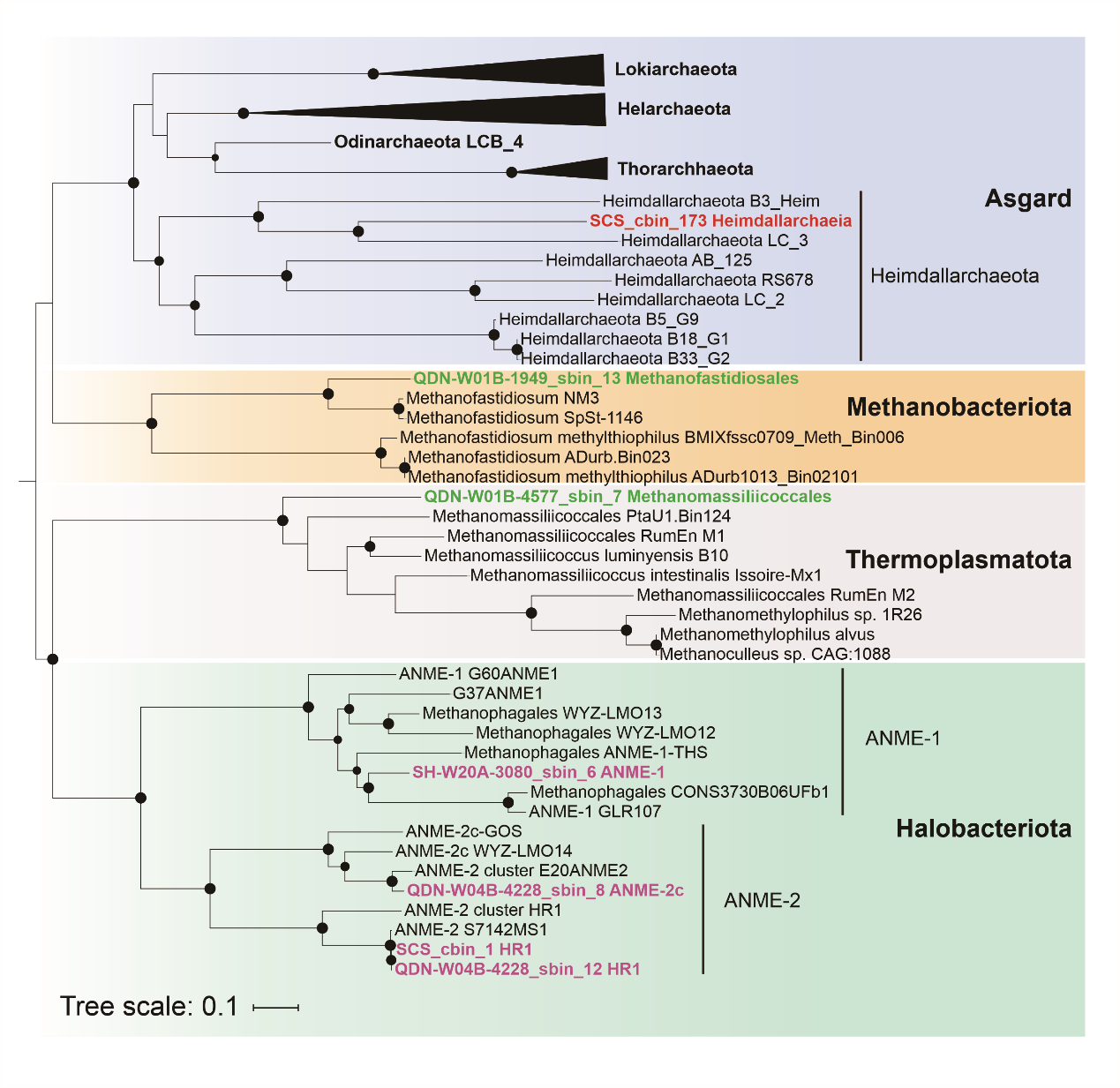

**Figure S10 Phylogenomic placement of the MAGs based on 43 conserved protein sequences.** Black dots indicate bootstrap values of 70–100%. Scale bars indicate the average number of substitutions per site.

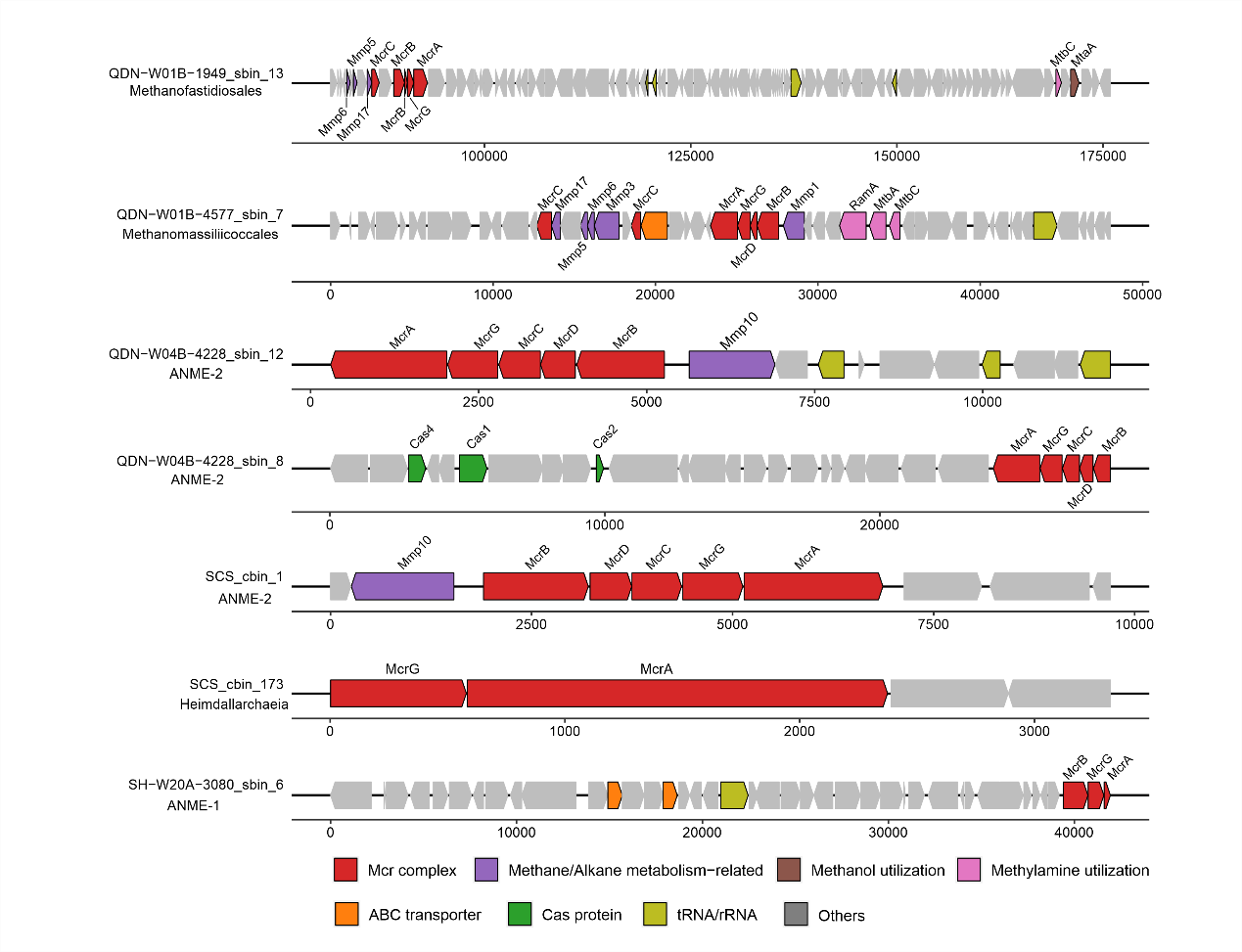

**Figure S11 Co-localization of *mcr* with genes nearby.** Genes are colored according to their functions. Red: Mcr complex; Purple: Methane/Alkane metabolism-related; Brown: Methanol utilization; Pink: Methylamine utilization; Orange: ABC transporter; Green: Cas protein; Greenyellow: tRNA/rRNA; Grey: Others.
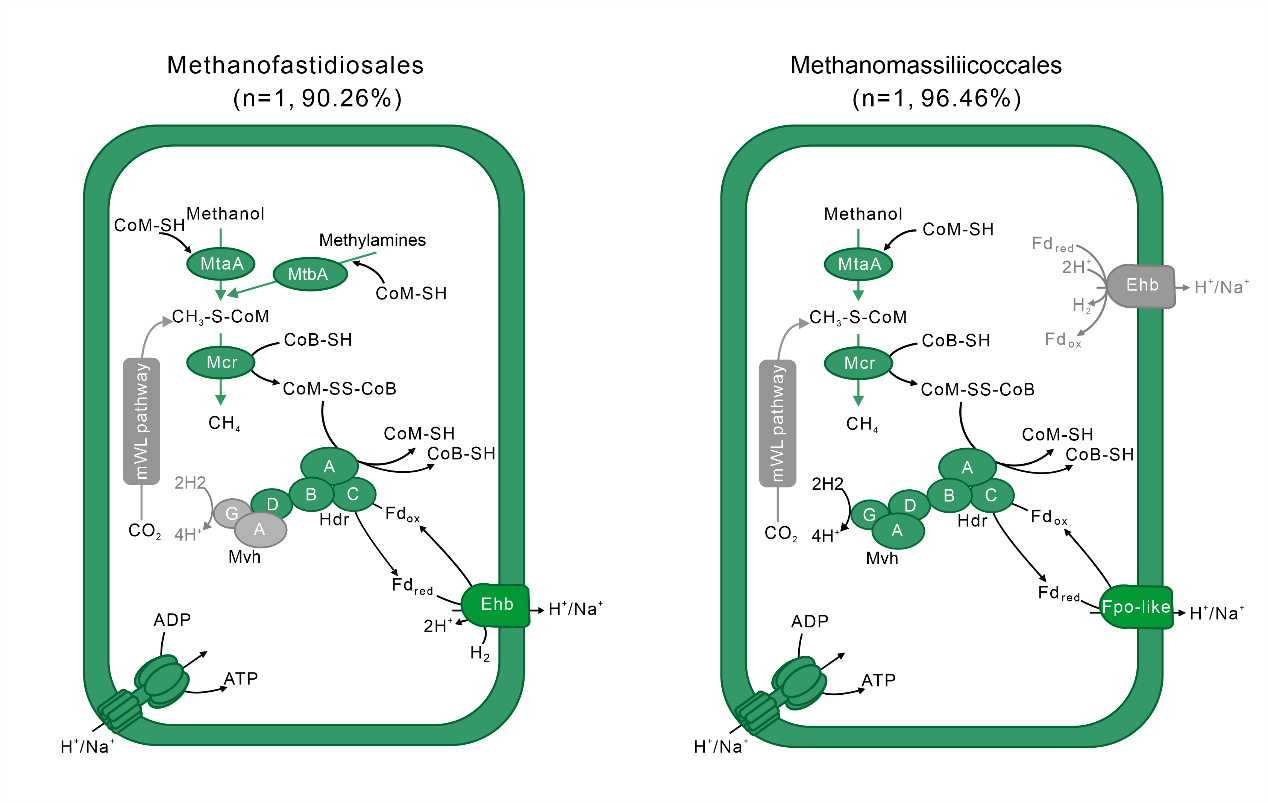

**Figure S12 Predicted pathways of hydrogen-dependent methylotrophic methanogenesis.** Grey colors indicate the absence of the enzyme or pathway. mWL pathway, methyl-branch of the Wood–Ljungdahl pathway. The percentages between brackets indicate the estimated completeness of the corresponding MAGs. A complete list of metabolic information can be found in Table S11.

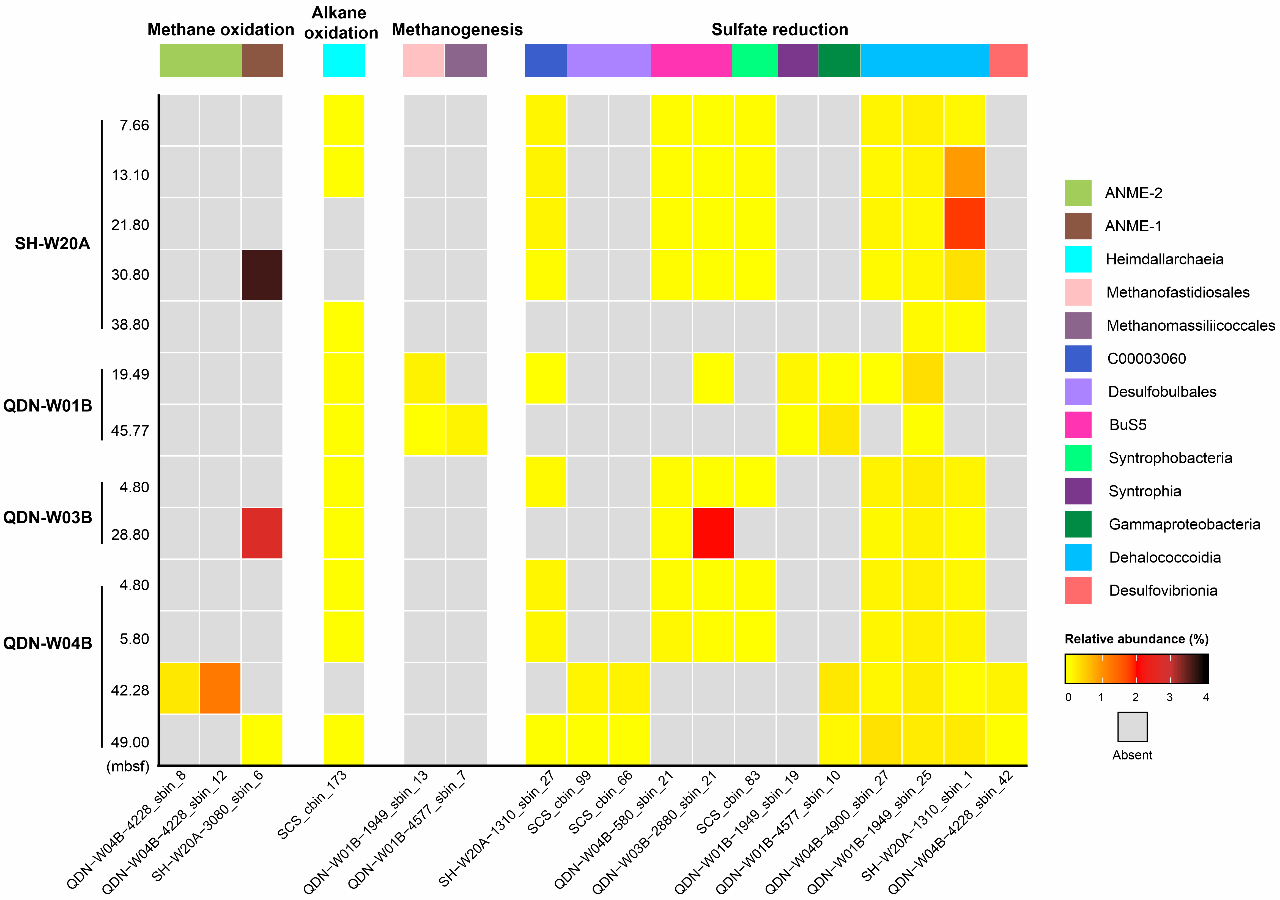

**Figure S13. Relative abundances of hydrocarbon-metabolizing archaea and sulfate reducers inferred from metabolic pathway reconstructions.** The MAG with the highest genome quality from each species cluster is picked as the representative. Details on species clusters and their relative abundances are presented in **Table S4**.

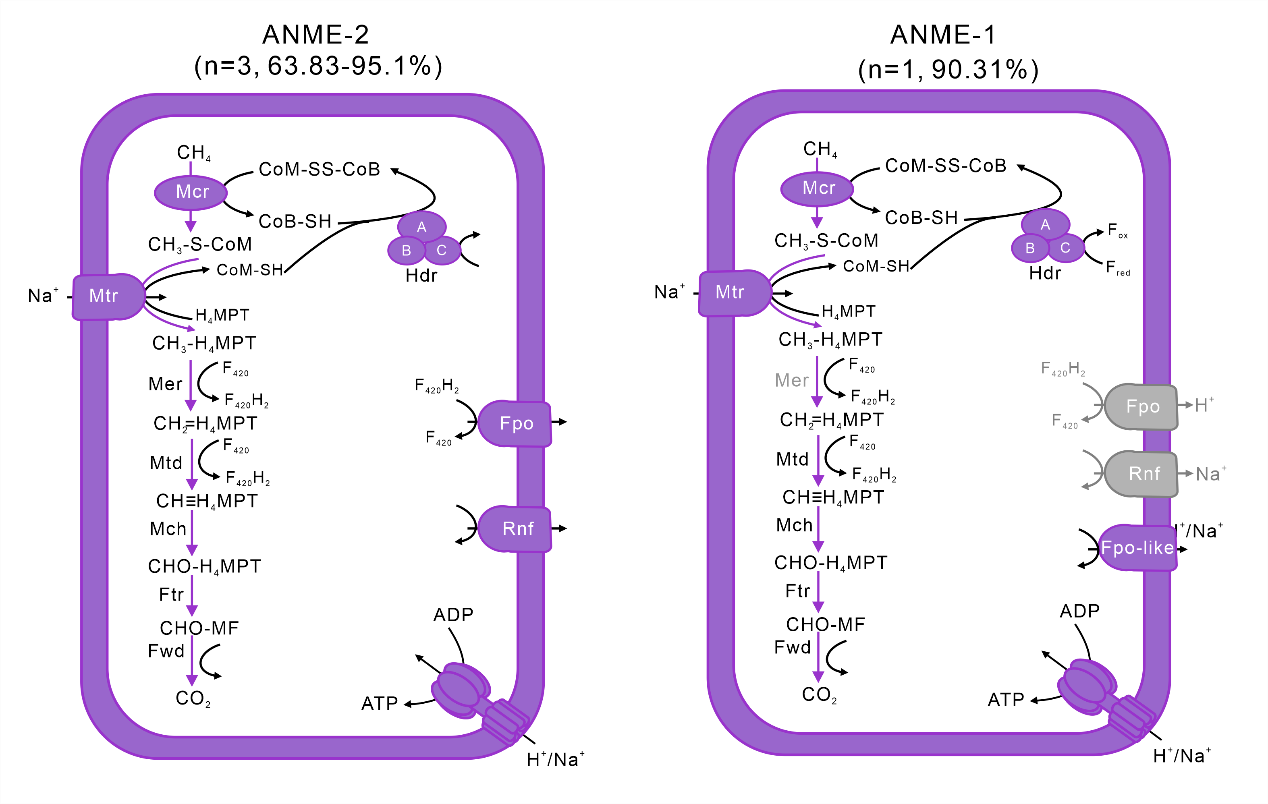

**Figure S14 Predicted pathways of Anaerobic oxidation of methane.** Grey colors indicate the absence of the enzyme or pathway. The percentages between brackets indicate the estimated completeness of the corresponding MAGs. A complete list of metabolic information can be found in Table S11.
